# “Glutaminolysis Fuels Reactive Astrocytes, Exacerbating Amyloid Pathology in Alzheimer’s Disease”

**DOI:** 10.1101/2025.06.12.658977

**Authors:** Shamulailatpam Shreedarshanee Devi, Sofia Luella Heese, Isabelle Leray, Gilbert Gallardo

## Abstract

Amyloid-β (Aβ) plaques with progressively increasing reactive astrocytes characterize Alzheimer’s disease (AD). Reactive astrocytes are regulated by cellular and molecular mechanisms that are known to progress Aβ pathology. However, the metabolic adaptation and metabolites required to fuel these molecular changes in reactive astrocytes remain unknown. Using human AD samples, *in vivo* amyloid mouse models, and *in vitro* approaches, we demonstrate that reactive astrocytes utilize glutamine to fuel anaplerosis and meet their metabolic demands, thereby progressing amyloidosis. We show that reactive astrocytes increase Na^+^-coupled neutral amino acid transporters for glutamine uptake that are interdependent on Na^+^/K^+^ ATPase. Furthermore, increasing brain-glutamine levels with a high-glutamine diet exacerbated reactive astrocytes, increasing Aβ burden in an amyloid mouse model. We demonstrate that glutamine undergoes glutaminolysis via glutaminase-2/glutamate dehydrogenase-1 enzymes to be incorporated into TCA metabolites for anaplerosis. Pharmacologically or genetically blocking glutaminolysis reduces reactive astrocytes and decreases Aβ pathology in an amyloid mouse. Our findings reveal the first glutamine-dependent metabolic adaptation of reactive astrocytes affecting Aβ pathology, which may be harnessed for AD therapeutic strategies.

## Introduction

Alzheimer’s disease (AD) is associated with progressively increased reactive astrocytes correlating with amyloid-β (Aβ) plaque pathology, neurofibrillary tau tangles, and cognitive decline. Reactive astrocytes are characterized by cellular changes including hypertrophic morphology, altered gene expression, and secretion of cytokines, suggesting that a highly metabolically dynamic cellular process is necessary for reactivity. Earlier studies have shown that reactive astrocytes in AD are associated with deviant metabolic demands, such as increased glucose and acetate uptake^1,2^, further indicating an altered metabolite obligation during reactivity. Nonetheless, the metabolic fate of glucose or acetate remains underdetermined, and whether they regulate reactive astrocytes in AD has not been shown. Therefore, the metabolic adaptation and the fate of any adaptive metabolites that fuel and regulate reactive astrocytes remain unknown.

Apart from glucose, glutamine is the most abundant metabolite in the mammalian system, with equal concentrations in cerebrospinal fluid (CSF) and plasma, facilitated by constant exchange across the blood-brain barrier (BBB). In the periphery, glutamine is a distinct amino acid that is metabolized through glutaminolysis for anaplerosis, a metabolic adaptation that replenishes the tricarboxylic acid (TCA) cycle metabolites. However, in the CNS, the presumed primary role of glutamine is to serve as the substrate for generating glutamate in the neurons that are supplied by glutamine-derived astrocytes. This concept, although rudimentary and oversimplified, has prevailed without considering that astrocytes express glutamine Na^+^-coupled neutral amino acid transporters (SNATs) and glutaminolysis enzymes, indicating that astrocytes can uptake glutamine and undergo glutaminolysis. Moreover, the cell-intrinsic role of glutamine and metabolic fate in sustaining the astrocytes’ reactivity in AD has not been explored yet.

Glutamine, a conditionally essential amino acid, is transported across the membrane by SNATs. Reports have suggested that the function of SNAT transporters is interdependent on Na^+^/K^+^ ATPase activity^3^. Previously, we have shown that reactive astrocytes in AD and amyotrophic lateral sclerosis (ALS) exhibit upregulation of the astrocytic isoform of α2-Na^+^/K^+^ ATPase (NKA)^4,5^. However, the cellular purpose of reactive astrocytes for increasing NKA activity and its downstream metabolic regulatory role during reactivity had not been explored. We hypothesize that an increase in NKA in reactive astrocytes might promote the SNAT transporter function for increased glutamine transport and glutaminolysis in reactive astrocytes. Here, we described the role of NKA/SNAT2-dependent glutamine uptake in reactive astrocytes required to fuel the TCA cycle for a reactive response for the first time. We also demonstrated that increasing glutamine delivery into the CSF with a high glutamine diet exacerbates reactive astrocytes that escalate Aβ-pathology in an AD mouse model for amyloidosis. We further demonstrate an in-depth understanding of glutamine metabolism through the actions of glutaminase (GLS) and glutamate dehydrogenase 1 (GLUD1), which serve as an anaplerotic pathway that fuels the TCA cycle, a process necessary for reactive astrocytes. Lastly, we developed a novel CRISPR/Cas9-based genetic tool for modifying and blocking the astrocytic glutaminolysis pathway, resulting in the suppression of reactive astrocytes, thereby decreasing amyloidosis in the AD amyloid mouse model. In summary, we identify a previously unknown astrocytic glutamine-dependent metabolic adaptation fueling reactivity, revealing several metabolites and enzymes, providing tools and targets for potential AD therapeutic strategies

## Result

### Astrocytic NKA knockdown decreased reactive astrocytes and Aβ pathology

To further investigate the NKA’s role in reactive astrocytes under pathological conditions of amyloidosis, we bred NKA conditional knockout mice (NKA^flox/flox^) with the APPswe/PS1d9 (APP/PS1) double transgenic mouse model of amyloidosis (APP/PS1-NKA^−/flox^)^6^. In response to Aβ plaques, reactive astrocytes surround plaques in human AD pathology, exhibiting increased expression of reactive markers glial fibrillary acidic protein (GFAP) and vimentin (VIM),and S100 calcium-binding protein β (S100β)^7–9^. This astrocytic response to Aβ plaques is also evident in APP/PS1 mice and is known to progress Aβ pathologies^10–12^. The APP/PS1^NKA-/flox^ mice were aged 8 months, followed by the NKA astrocytic conditional knockout via Adeno-associated virus-mediated (AAV) expression of Cre. We unilaterally injected AAVs encoding Cre under the GFAP promoter or control AAVs into the hippocampus of 8-month-old APP/PS1^NKA-/flox^ mice, a time point after disease onset. We and others have previously utilized viral-mediated strategies for astrocytic gene knockout studies, including AAVs^4,5,13–16^. Following Cre expression, the ipsilateral hippocampus exhibited decreased NKA expression four months post-injection, determined by immunofluorescence (IF) (Figure S1A). At 12 months, we analyzed reactive astrocytes by IF, revealing that astrocytic NKA knockdown decreased GFAP and VIM surrounding fibrillary amyloid plaques (X34) relative to control APP/PS1 mice (Figures 1B, 1C, 1D, 1E). Notably, decreasing astrogliosis significantly reduced total Aβ plaques (6E10) and fibrillary plaques (X34) following astrocytic NKA knockdown in APP/PS1^NKA-/flox^ mice compared to the control APP/PS1 mice (Figures 1D, 1F, 1G, 1H). A pathological feature of AD is the presence of swollen and bulbous-shaped degenerating dystrophic neurites that accumulate lysosomal-associated membrane protein 1 (LAMP-1) around plaques^17^. Additional analysis of LAMP1 revealed that astrocytic NKA knockdown preserved neurons, decreasing dystrophic neurites compared to control APP/PS1 mice (Figures 1I, 1J). Our data show that conditional astrocytic NKA decreases reactive astrocytes, Aβ plaques, and dystrophic neuritis.

**Figure 1.**
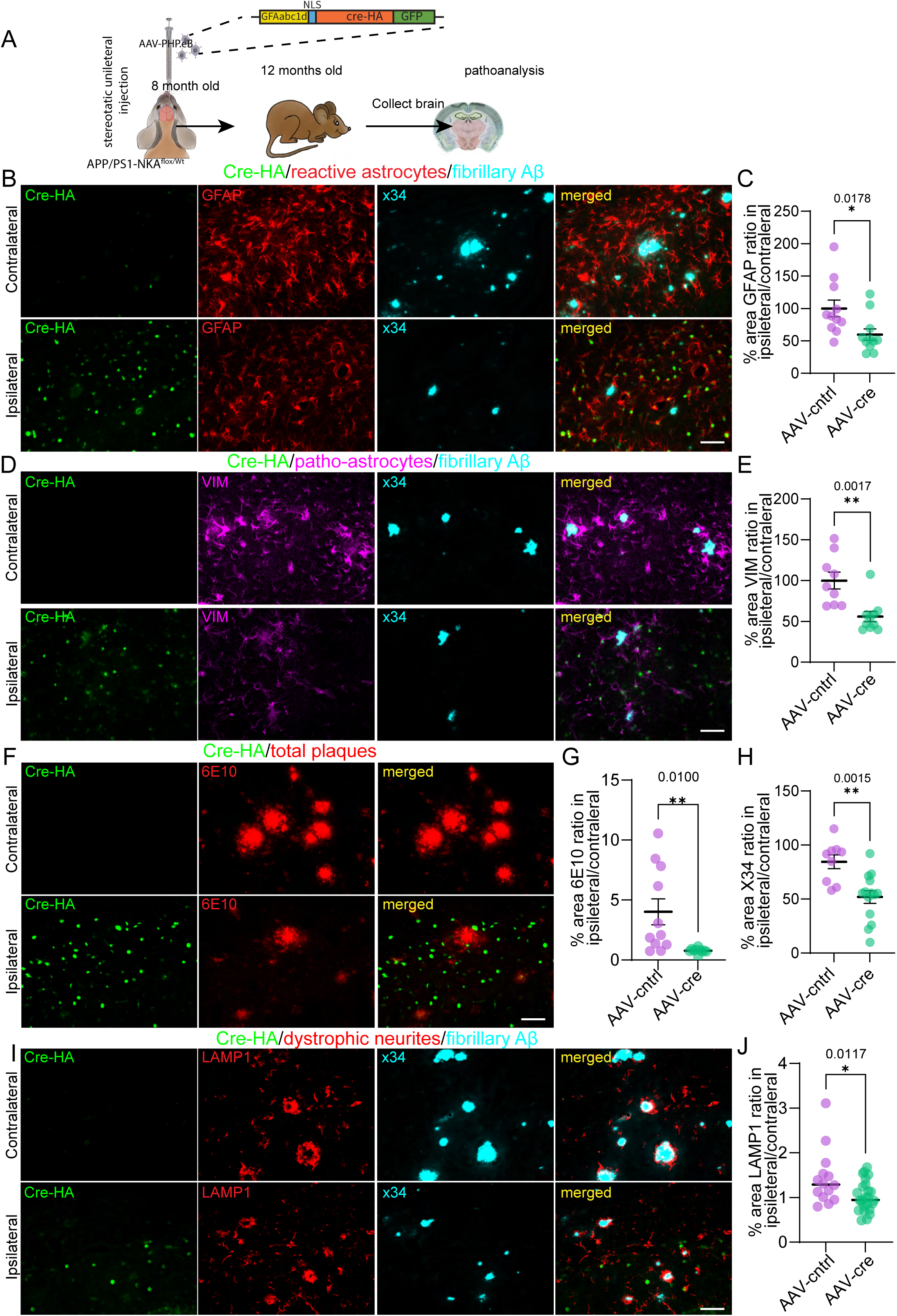
Astrocytic NKA knockdown decreased reactive astrocytes and Aβ pathology. (A) Schematic for the timing of AAV-GFAP-Cre injections into female/male APP/PS1 mice. (B) IF for Cre-HA, astrocytes, and fibrillary plaque in the hippocampus of APP/PS1-NKA knockdown and control. Scale bar, 30μm. (C) Quantification of the percent area of astrocytes in APP/PS1-NKA knockdown and control. n=6-8. (D) IF for Cre-HA, astrocytes, and fibrillary plaque in the hippocampus of APP/PS1-NKA knockdown and control. Scale bar, 30μm. (E) Quantification of the percent area of astrocytes in APP/PS1-NKA knockdown and control. n=6-8 (F) IF for Cre-HA and total Aβ plaque in the hippocampus of APP/PS1-NKA knockdown and control. Scale bar, 30μm. (G) Quantification of the percent area covered by total Aβ plaques in APP/PS1-NKA knockdown and control. n=6-8(H) Quantification of the percent area covered by fibrillary plaques in APP/PS1-NKA knockdown and control. n=6-8 (G) IF for Cre-HA, dystrophic neurites, and fibrillary plaque in the hippocampus of APP/PS1-NKA knockdown and control. Scale bar, 30μm.(H) Quantification of the percent area covered by dystrophic neurites in APP/PS1-NKA knockdown and control. n=6-8. Significance determined by two-tailed unpaired student *t* test. Error bars represent ± sem. *p<0.5, **<0.005.

### Reactive astrocytes increase glutamine uptake in an NKA-SNAT2 interdependent mechanism

We next utilized primary astrocytes to investigate the NKA’s mechanistic insights by inducing a proinflammatory response with a cocktail of chemokines/cytokines (TNF-α, IL-1β, and IL-6). Studies have illustrated those reactive astrocytes in pathological conditions, including AD, are nuclear factor kappa B (NF-ƙB) transcription factor-dependent, leading to proinflammatory gene expression, promoting disease progression^10,12^. Therefore, we evaluated NF-ƙB activity in primary astrocytes following cyto/chemo activation with a lentiviral-mediated reporter encoding GFP under the NF-ƙB response element promoter (Figure 2A). The NF-ƙB reporter assay showed that primary astrocytes stimulated with chemokines/cytokines (Ast^C+C^) significantly increased GFP expression compared to control after 48hrs (Figure 2B, 2C). Next, we analyzed the expression of several proinflammatory genes in reactive Ast^C+C^ by qPCR, revealing a significant proinflammatory response after 48hrs (Figure S1B). This data illustrates that our primary reactive Ast^C+^ is an NF-κB-dependent proinflammatory response, mimicking the reactive astrocytes observed in AD, enabling the investigation of NKA’s mechanistic insights.

**Figure 2.**
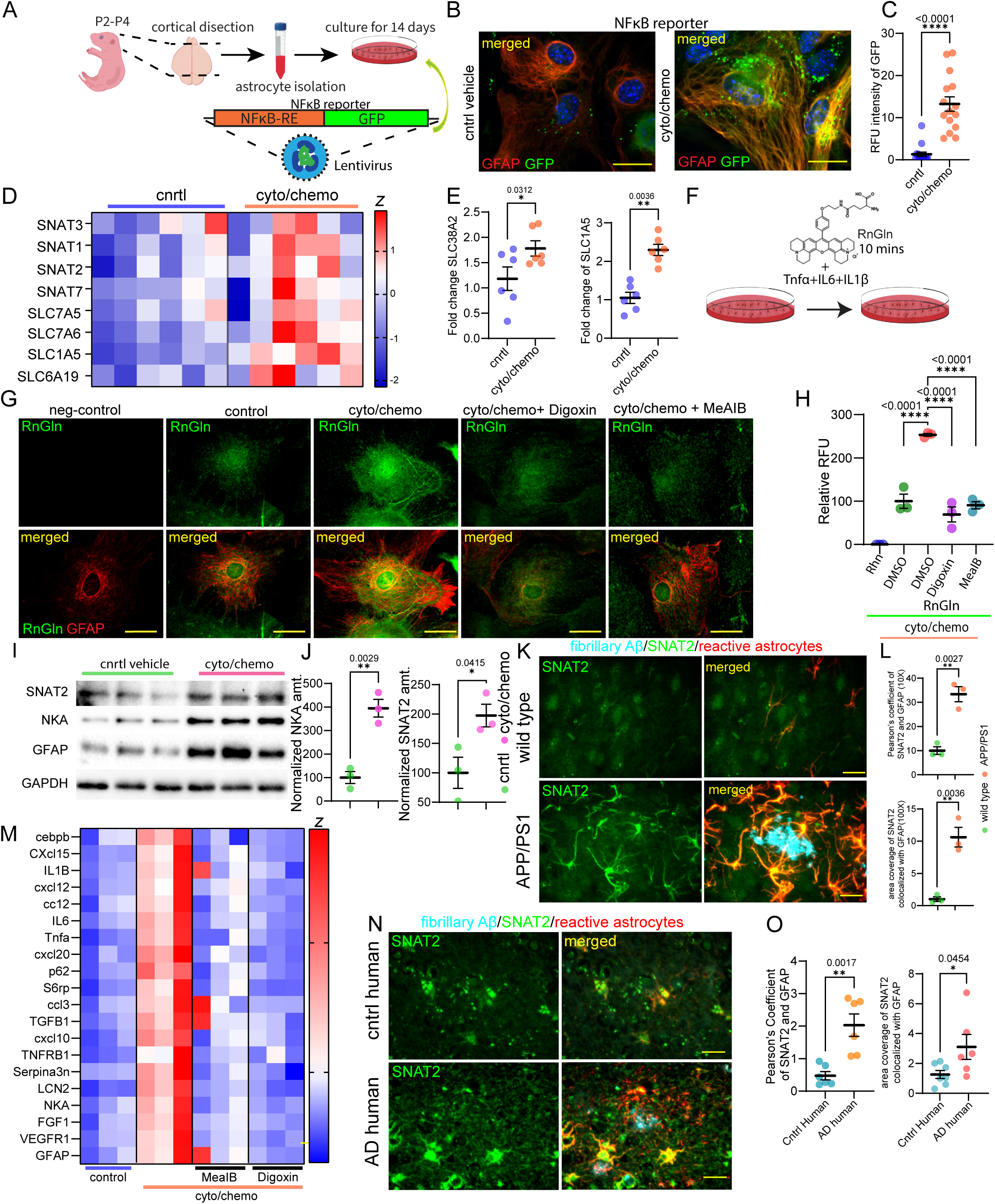
Reactive astrocytes increase glutamine uptake in an NKA-SNAT2 interdependent mechanism. (A) Schematic for primary astrocytes isolation and lentivirus transduction encoding NFκB-GFP reporter. (B) IF of primary astrocytes expressing NFκB-GFP reporter following cyto/chemo activation. (C) Quantification of relative fluorescent units (RFU) for NFκB-GFP activity in reactive primary astrocytes compared to control. n=3; Scale bars, 10 μm (D) Gene expression heat map for SNATs in reactive primary astrocytes and control. Z score; n=6. (E) Quantification gene expression fold change for *SLC38A2* and *SLC1A5* in reactive primary astrocytes compared to control normalized to GAPDH. n=6 (F) Schematic depicting rhodamine-labeled glutamine uptake assay in primary astrocytes. (G) IF for Rho-Glutamine in primary astrocytes, reactive astrocytes, and NKA or SNAT inhibition colocalized with GFAP. Scale bars 10μm (H) Quantification of Rho-Glutamine RFU in control primary astrocytes, reactive astrocytes, and NKA or SNATs inhibition. n=3 (I) Immunoblotting for SNAT2, NKA, and GFAP protein levels in reactive primary astrocytes and control. n=3. (J) Densitometric quantification of SNAT2 and NKA protein levels in reactive primary astrocytes relative to control normalized to GAPDH. n=3 (K) IF for SNAT2, astrocytes, and fibrillary plaques in APP/PS1 and control wild-type mice. Scale bars 30μm (L) Quantification of the percentage of SNAT2 colocalized with astrocytes around plaques in APP/PS1 mice compared to control wild-type mice (top graph). Pearson’s coefficient of SNAT2 and GFAP (bottom graph), n=3 (M) Heat map of proinflammatory gene expression in reactive astrocytes and SNATs or NAK inhibition compared to control. Z score; n=6 (N) IF for SNAT2, astrocytes, and fibrillary plaques in human AD brains and control humans. Scale bars 30μm(O) Quantification of the percentage of SNAT2 colocalized with GFAP around plaques in human AD brains compared to control humans (top graph). Pearson’s coefficient of SNAT2 and GFAP (bottom graph), n=6-7 two brain sections per human subject. Data significance was calculated using a two-tailed unpaired Student t-test or a one-way ANOVA with Bonferroni post-hoc tests when appropriate. Error bars represent ± sem. *p<0.5, **<0.005, ***p<0.0005, ****p<0.0001

The previously reported interplay between SNATs and the NKA raised the question of whether glutamine transport has a role in reactive astrocytes. To address this notion, we evaluated the expression of Na^+^-dependent SNAT transporters in reactive Ast^C+C^, identifying several upregulated influx transporters, SNAT1, SNAT2, SNAT7, and SLC6A19, together with bidirectional transporters ASCT2, SNAT3, SLC7A5, and SLC7A6 (Figure 2D, 2E). The increased expression of both influx SNATs and bidirectional glutamine transporters in reactive astrocytes led us to investigate the direction of glutamine flow by using rhodamine-labeled glutamine (RnGln)^18^ treated conditioned media. We found that reactive Ast^C+C^ significantly increased RnGln import into the cytosol compared to the control vehicle (Figure 2F, 2G, 2H). We next evaluated the NKA’s role in increasing glutamine import by blocking NKA activity with Digoxin, which significantly decreased RnGln import in reactive Ast^C+C^ (Figure 2G, 2H). We further inhibited SNATs with α-(methylamino) isobutyric acid (MeaIB), which also significantly decreased RnGln import in reactive Ast^C+C^ (Figure 2G, 2H). The increase in glutamine in reactive Ast^C+C^ also correlated with increased SNAT2 and NKA expression, determined by immunoblotting analysis (Figure 2I, 2J). Next, we determined if increased glutamine import is necessary for a proinflammatory response in reactive Ast^C+C^. To test this notion, we inhibited SNATs or the NKA, blocking glutamine import in reactive Ast^C+C^, and analyzed proinflammatory gene expression by qPCR. This analysis revealed that blocking glutamine import suppressed the expression of several proinflammatory genes in chemokines/cytokines stimulated Ast^C+C^ (Figure 2M). These findings support our hypothesis that reactive astrocytes increase glutamine import, which is interdependent on the NKA and SNAT transporters. Moreover, our findings suggest that SNAT-dependent glutamine import is essential for reactive astrocytes’ proinflammatory response.

To evaluate whether reactive astrocytes also increase glutamine SNAT transporters, we next analyzed 18-month-old APP/PS1 mice displaying measurable Aβ plaques, revealing significantly increased SNAT2 expression colocalized with astrocytes (GFAP) around plaques (X34) (Figure 2K, 2L). However, SNAT2 levels colocalizing with neuronal marker MAP2 were not altered in APP/PS1 mice (Figure S2A, S2B). We further validated that the SNAT2 glutamine transporter is significantly increased in reactive astrocytes in human AD brain subjects compared to age-matched human controls, determined by IF analyses (Figure 2N, 2O). These findings suggest that astrocytes increase SNAT2 glutamine transporter expression in response to pathological stimuli and may be a key astrocyte pathological feature in AD pathophysiology. In addition, the increased SNAT2 expression in reactive astrocytes suggests that glutamine import may regulate their proinflammatory response. On the contrary, SNAT1 was not expressed in astrocytes in APP/PS1 mice and was found to be predominantly neuronal, as previously reported (Figure S2C, S2D)^19^.

### Increasing CNS-glutamine levels exacerbate reactive astrocytes *in vitro and in vivo* in APP/PS1 mice

Our findings that reactive astrocytes increase SNAT2 and glutamine import suggested that higher glutamine concentrations may also further influence their reactivity. Therefore, we evaluated reactive Ast^C+C^ proinflammatory response upon increasing glutamine levels in the conditioned media to 2mM, 4 times higher than the physiological 0.5 mM concentration. Surprisingly, high-glutamine-treated Ast^C+C^ (high-Gln) further increased proinflammatory gene expressions compared to physiological 0.5mM concentrations, suggesting increasing glutamine availability exacerbates the astrocytic proinflammatory response (Figure S3A).

Glutamine is a conditional amino acid for which biosynthesis alone is insufficient for cellular function, making it reliant on glutamine import from the diet^20^. As glutamine import increases in reactive primary astrocytes and higher glutamine concentrations exacerbate their proinflammatory response, we hypothesized that increasing CNS-glutamine levels would further aggravate astrogliosis. Glutamine from the diet is readily absorbed, resulting in a transient increase in plasma levels^21,22^. Therefore, we first tested whether increasing dietary glutamine could increase CNS glutamine levels by providing a high-glutamine diet daily to a cohort of 6-month-old female/male mice. Analyzing the CSF following a daily high-glutamine diet revealed a significant increase in glutamine levels in female/male mice compared to the control diet, enabling us to test our hypothesis that increasing CNS-glutamine concentrations aggravates reactive astrocytes (Figure 3B). Although female mice displayed a significant increase in plasma glutamine levels, no significant difference was found in males following a high-glutamine diet, potentially due to the higher muscle mass that readily absorbs glutamine (Figure 3B).

**Figure 3.**
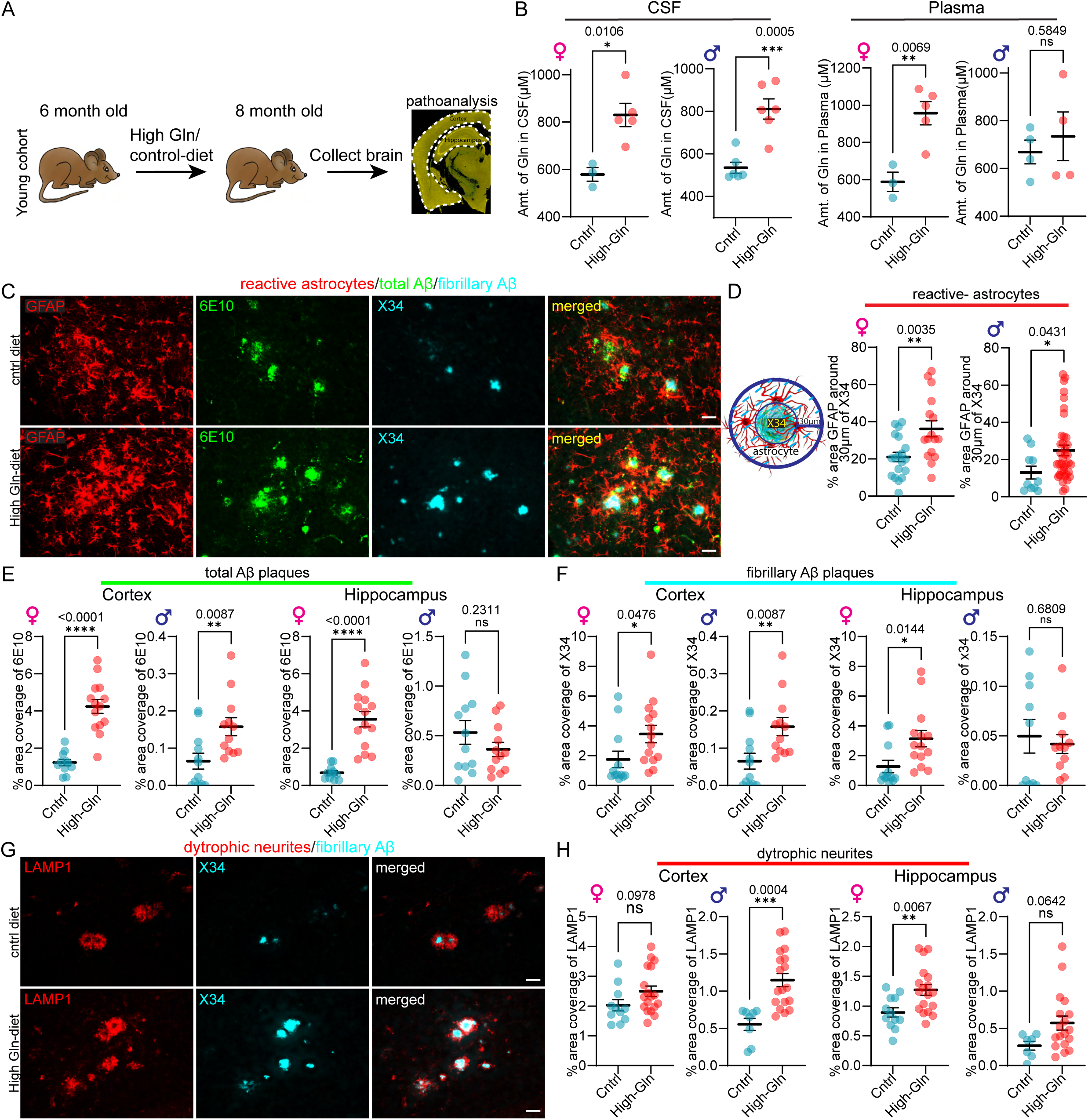
Increasing CNS-glutamine levels exacerbate reactive astrocytes *in vitro and in vivo*. (A) Schematic of a high-glutamine diet timeline before overt amyloidosis in APP/PS1 mice. (B) Quantification of CSF and plasma glutamine levels following a high-glutamine diet compared to control diet. n=3-6 (C) IF for astrocytes, total plaques, and fibrillary plaques following a high-glutamine diet or control diet in APP/PS1 mice. Scale bars 30μm (D) Quantification of the percent area of astrocytes around a 30-μm radius of fibrillary plaques following a high-glutamine diet compared to control diet. n=5-7(E) Quantification of the percent area covered by total plaques in the cortex/hippocampus following a high-glutamine diet compared to control diet. n=5-7 (F) Quantification of the percent area covered by fibrillary plaques in the cortex/hippocampus following a high-glutamine diet compared to control diet. n= 5-7 (G) IF for dystrophic neurites and fibrillary plaques following a high-glutamine diet or control diet in APP/PS1 mice. Scale bars 30μm (H) Quantification of the percent area covered by dystrophic neurites in the cortex/hippocampus following a high-glutamine diet compared to control diet. n= 5-7. Data significance was calculated using a two-tailed unpaired Student t-test. Error bars represent ± sem. *p<0.5, **<0.005, ***p<0.0005, ****p<0.0001

To test our hypothesis, we fed an aged cohort of female/male APP/PS1 mice a high-glutamine diet starting at 6 months before overt Aβ pathology (APP/PS1^HG_Young^). The APP/PS1 mice develop Aβ pathology earlier in female mice, ∼6 months, while the male mice start closer to 8 months. Following two months of a high-glutamine diet at 8 months, APP/PS1^HG_Young^ mice showed an increase in reactive astrocytes in both female and male mice compared to their littermate APP/PS1 mice on a control diet, as measured by IF analysis of GFAP-positive astrocytes around 30µm of fibrillary plaques (X34) (Figure 3C, 3D). This data corroborates our results on primary astrocytes, which showed increased activity upon increasing glutamine concentrations.

We next analyzed total Aβ plaque burden (6E10) following high-dietary glutamine in APP/PS1^HG_Young^, finding a significant increase in both females and males in the cortex compared to the control diet, consistent with an increase in reactive astrocytes (Figures 3C, 3E). In the hippocampus, we found a significant increase in total Aβ plaques in females with no substantial change in male APP/PS1^HG_Young^ mice compared to littermate APP/PS1 mice on a control diet (Figure 3E). We next evaluated fibrillary plaques (X34) that are associated with neurodegeneration. Consistent with the total plaques in the cortex, fibrillary plaque levels were significantly increased in both female and male APP/PS1^HG_Young^ compared to the control diet (Figures 3C, 3F). We further found fibrillary plaques were also significantly increased in the hippocampus of female APP/PS1^HG_Young^ with no substantial change in males APP/PS1^HG_Young^ compared to the control diet (Figure 3F). We next analyzed the effect of increasing CNS glutamine levels on dystrophic neurites (LAMP1), finding a significant increase in male cortex with a trending increase in female APP/PS1^HG_Young^ compared to the control diet (Figures 3G, 3H). In the hippocampus, female APP/PS1^HG_Young^ displays a significant increase in dystrophic neurites with a trending increase in males (Figure 3H). These analyses demonstrate that increasing CNS glutamine levels early in amyloidosis exacerbates astrogliosis, Aβ pathologies, and dystrophic neurites.

Previously, we and others reported that altering the astrocytic responses further regulates microglia in mouse models of AD pathology^5,23^. Therefore, we evaluated microgliosis in APP/PS1^HG_Young^, finding a gender-dependent difference in the cortex with a significant increase in microgliosis marker IBA1 in males and a considerable decrease in females compared to the control diet (Figure S3C, S3D). In the hippocampus regions, the microglial response was unaltered in both APP/PS1^HG_Young^ genders, suggesting that the increased Aβ pathologies largely depend on increased reactive astrocytes in early disease (Figure S3D). We also analyzed the weight change throughout the high-glutamine diet in APP/PS1^HG_Young^ mice, as previous studies reported that glutamine supplementation induces weight loss in animal studies. Following an initial weight drop after a high-glutamine diet, female and male APP/PS1^HG_Young^ mice recovered weight gain faster than the control (Figure S3E). However, contrary to previous reports, 8 weeks of a high glutamine diet led to no significant changes in overall weight in APP/PS1^HG_Young^ mice.

### Increasing CNS-glutamine levels after disease onset exacerbates reactive astrocytes and Aβ pathologies

AD’s pathology develops in the early decades before measurable symptoms and clinical diagnosis. Therefore, it is essential to determine whether increasing CNS-glutamine levels after disease onset and pronounced Aβ pathology further exacerbate reactive astrocytes and amyloidosis. To evaluate this notion, we fed a high-glutamine diet to a cohort of female/male APP/PS1 starting at 12 months after disease onset (APP/PS1^HG_Old^) for two months. Similar to the younger cohort, the APP/PS1^HG_Old^ mice regained weight much faster in both genders compared to the control treatment (Figure S4B). At 14 months, APP/PS1 mice have measurable Aβ pathologies in various brain regions. Thus, we quantified pathologies in the hippocampal regions, specifically coronal-1 (CA1), coronal-3 (CA3), and dentate gyrus (DG), as well as the piriform cortex (PC). The IF analysis of GFAP surrounding plaques following a high-glutamine diet revealed a significant increase in reactive astrocytes in the hippocampal CA1/DG and PC with a strong trending increase in all other brain regions of female APP/PS1^HG_Old^ mice compared to the control diet (Figures 4B, 4C, and S4D). The analysis of males APP/PS1^HG_Old^ mice displayed a significant increase in reactive astrocytes in the hippocampal DG and PC, with a strong trending increase in all other brain regions compared to the control diet (Figures 4C, and S4D).

**Figure 4.**
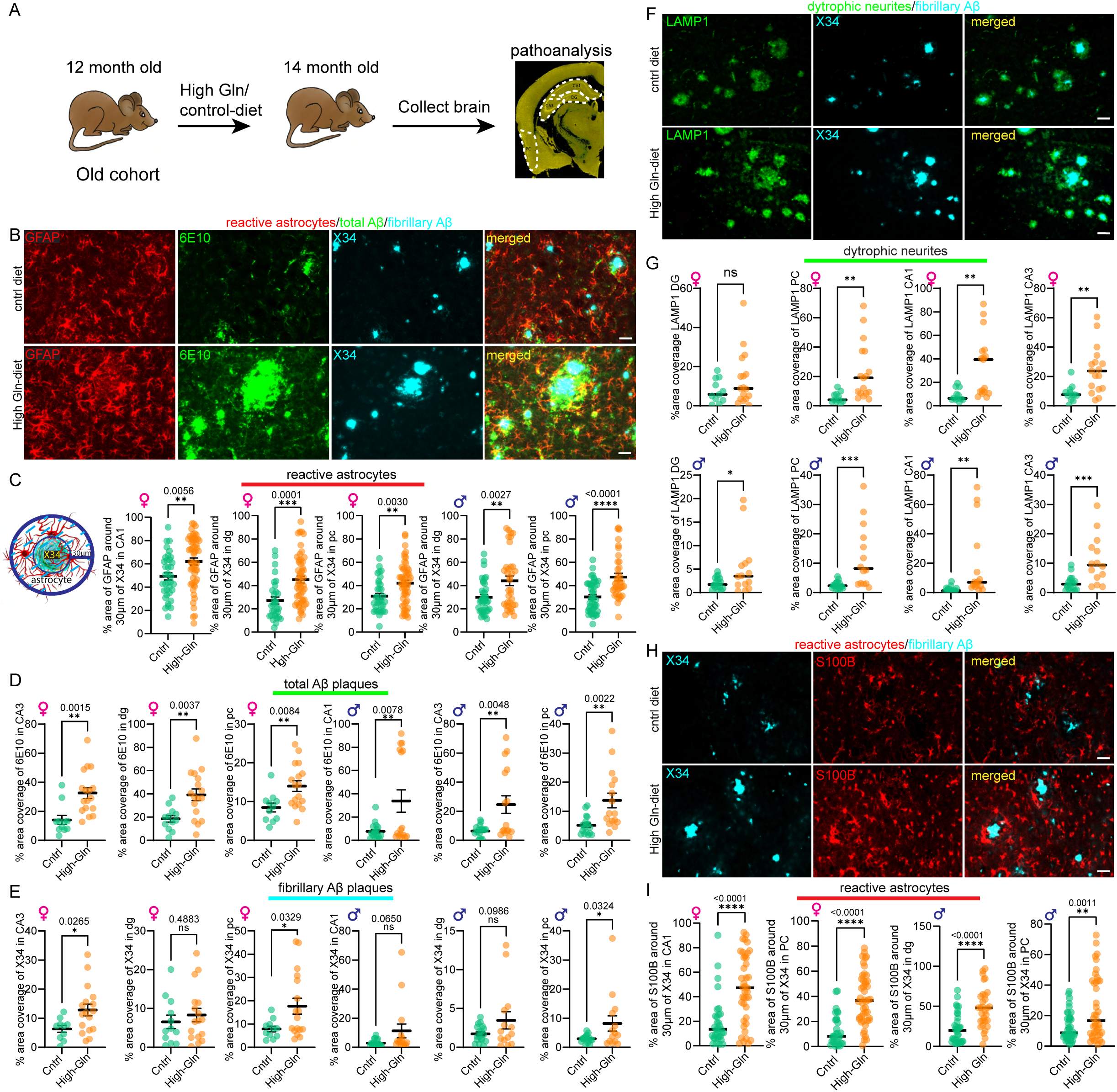
Increasing CNS-glutamine levels after disease onset exacerbates reactive astrocytes and Aβ pathologies. (A) Schematic diagram of a high-glutamine diet timeline after disease onset in APP/PS1 mice. (B) IF for astrocytes, total plaques, and fibrillary plaques following a high-glutamine diet or control diet in APP/PS1 mice. Scale bars 30μm (C) Quantification of the percent area of astrocytes around a 30-μm radius of fibrillary plaques following a high-glutamine diet compared to control diet. n= 6-9 (D) Quantification of the percent area covered by total plaques in the cortex/hippocampus following a high-glutamine diet compared to control diet. n= 6-9 (E) Quantification of the percent area covered by fibrillary plaques in the cortex/hippocampus following a high-glutamine diet compared to control diet. n= 6-9 (F) IF for dystrophic neurites and fibrillary plaques following a high-glutamine diet or control diet in APP/PS1 mice. Scale bars 30μm (G) Quantification of the percent area covered by dystrophic neurites in the cortex/hippocampus following a high-glutamine diet compared to control diet. n= 6-9(H) IF for astrocytes, total Aβ plaques, and fibrillary plaques following a high-glutamine diet or control diet in APP/PS1 mice. Scale bars 30μm (I) Quantification for percent area of astrocytes around a 30-μm radius of fibrillary plaques following a high-glutamine diet compared to control diet. n= 6-9 Data significance was calculated using a two-tailed unpaired Student t-test. Error bars represent ± sem. *p<0.5, **<0.005, ***p<0.0005, ****p<0.0001

We next analyzed total Aβ plaques (6E10), as previous studies have illustrated that pathologically reactive astrocytes increase linearly with plaque burden^24–27^. This analysis revealed a significant increase in total plaque in the PC and hippocampal CA3/DG, with a trending increase in the CA1 in female APP/PS1^HG_Old^ compared to the control diet (Figures 4B, 4D, and S4E). Male APP/PS1^HG_Old^ significantly increased total plaques in all brain regions compared to the control diet (Figures 4B, 4D, and S4E). We further evaluated fibrillar plaques (X34) and observed a significant increase in the PC in both APP/PS1^HG_Old^ genders and female hippocampal CA3 region, with all other brain regions displaying a strong trending increase in both genders compared to the control diet (Figures 4B, 4E, and S4F). Next, we analyzed LAMP1 by IF, finding male APP/PS1^HG_Old^ mice displayed significantly increased dystrophic neurites in DG, CA1 and CA3 in females and CA1, CA3 and PC in male with a strong trend in PC and DG in female and male respectively compared to the control diet (Figures 4F, 4G).

We further quantified S100β, a second astrocytic marker that increases linearly with corresponding amyloid pathology levels, finding a significant increase in all the hippocampal regions and PC in both genders of APP/PS1^HG_Old^ compared to the control diet (Figures 4H, 4I, and S4G). The increase in astrocytic s100β in APP/PS1^HG_Old^ mice suggests that the effect of high-glutamine is the phenotype of reactive astrocytes and is not solely dependent on GFAP-expressing astrocytes. To evaluate microglia, we analyzed IBA1 by IF. Surprisingly, we observed a significant decrease in the IBA1-positive microglia in the cortex and the hippocampus of APP/PS1^HG_Old^ mice in both genders (Figure S5A, S5B).

We further evaluated whether increasing CNS-glutamine levels potentially increased synaptic activity, which may increase Aβ-peptide production by IF analysis of neuronal c-Fos, a marker for neuronal activity^28,29^. The analysis of neuronal c-Fos showed no significant changes in the cortex or hippocampus in both APP/PS1^HG_Old^ genders compared to the control diet (Figure S5C, S5D). This suggests that the effect on amyloid pathology following a high glutamine diet is not necessarily due to increased neuronal activity. The consistently increased astrocytic response suggests that the astrocytes are critical in exacerbating amyloidosis in high CNS-glutamine conditions after disease onset.

### Reactive astrocytes increase glutaminolysis enzymes in APP/P1 mice and human AD brains

To better understand the mechanistic insights into glutamine metabolism in reactive astrocytes, we hypothesized that reactive astrocytes increase glutamine import for glutaminolysis, which is necessary for a proinflammatory response. To test our hypothesis, we analyzed GLS2 protein levels, which catalyze glutamine’s conversion to glutamate in aged APP/PS1 mice. The IF analysis demonstrated that GLS2 expression was significantly increased in reactive astrocytes (GFAP) surrounding fibrillary plaques in APP/PS1 mice compared to control wild-type mice (Figure 5A, 5B). We further validated in human AD brain sections that GLS2 was significantly increased in reactive astrocytes around Aβ plaques compared to human control brain samples (Figure 5C, 5D). We next analyzed GLUD1, the second enzyme that metabolizes glutamate to α-ketoglutarate (α-KG), a rate-limiting substrate for the TCA cycle. This analysis showed a significantly increased GLUD1 expression colocalizing in reactive astrocytes surrounding fibrillary plaques in aged APP/PS1 mice compared to controls (Figure 5E, 5F). Additionally, human AD brain sections also demonstrated that reactive astrocytes increase GLUD1 expression around Aβ plaques compared to control human samples (Figure 5G, 5H). These analyses suggested that during astrogliosis glutamine import is increased for glutaminolysis, potentially regulating the pathological response. However, GLUD1 analysis in neurons revealed no significant changes colocalized with neuronal marker MAP2 in APP/PS1 mice (Figure S6A).

**Figure 5.**
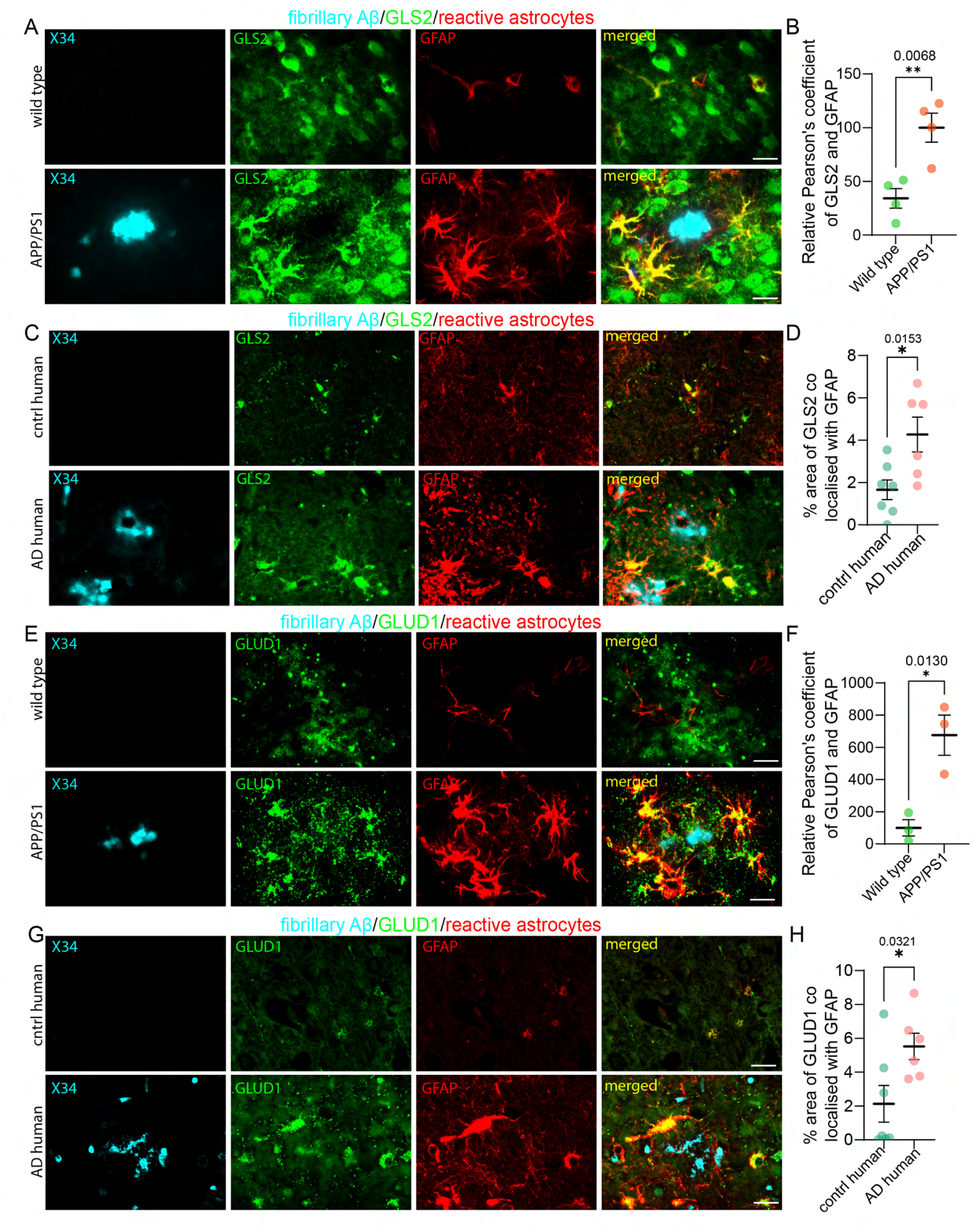
Reactive astrocytes increase glutaminolysis enzymes in APP/PS1 mice and human AD brains. (A) IF for GLS2, astrocytes, and fibrillary plaques in APP/PS1 and wild-type mice. Scale bars 30μm (B) Quantification of Pearson’s coefficient for GLS2 colocalized with astrocytes around plaques in APP/PS1 mice compared to wild-type mice. n=3 (C) IF for GLS2, astrocytes, and fibrillary plaques in human AD brains subjects and human controls. Scale bars 30μm (D) Quantification of Pearson’s coefficient for GLS2 colocalized with astrocytes around plaques in human AD brains compared to human controls. n=6-7 two brain sections per human subject. (E) IF for GLUD1, astrocytes, and fibrillary plaques in APP/PS1 and wild-type mice. Scale bars 30μm (F) Quantification of Pearson’s coefficient for GLUD1 colocalized with astrocytes around plaques in APP/PS1 mice compared to wild-type mice. n=3 (G) IF for GLUD1, astrocytes, fibrillary plaques in human AD brains and human controls. Scale bars 30μm (H) Quantification of Pearson’s coefficient for GLUD1 colocalized with astrocytes around plaques in human AD brains compared to human controls. n=6-7Data significance was calculated using a two-tailed unpaired Student t-test. Error bars represent ± sem. *p<0.5, **<0.005

### Glutaminolysis fuels reactive astrocytes for a proinflammatory response

The increased glutaminolysis enzymes in astrocytes of human AD brains and APP/PS1 mice suggested that glutamine metabolism may be critical in regulating an astrocytic pathological response. To comprehensively understand glutamine’s metabolism in reactive Ast^C+C^, we performed isotope tracing using U-^13^C_5_-glutamine. Strikingly, the abundance of ^13^C-glutamate and ^13^C-α-KG was significantly increased in reactive Ast^C+C^, relative to control, indicating an increase in glutaminolysis (glutamine→glutamate→αKG) by the enzymes GLS2/GLUD1 (Figure 6A). We next traced U-^13^C_5_-glutamine for TCA cycle metabolites, finding a significant increase in ^13^C incorporation in all metabolites compared to the control (Figure 6A). This data suggested that glutaminolysis serves as an anaplerotic reaction, replenishing the TCA metabolites. We further confirmed an increased TCA cycle by measuring ^13^C-labeled 2-hydroxyglutarate (2HG), a TCA cycle byproduct, finding a significant increase in reactive Ast^C+C^ compared to the control (Figure 6A). Next, we determined whether the increased proinflammatory response of reactive Ast^C+C^ in high-glutamine conditions further increased glutaminolysis for TCA cycle anaplerosis by treating Ast^C+C^ with a high 2mM U-^13^C_5_-glutamine in conditioned media. Isotope tracing analysis found high glutamine conditions increased ^13^C-glutamine fractional enrichment of glutamate and α-KG compared to reactive Ast^C+C^ in physiological 0.5mM glutamine concentrations (Figure 6A). In addition, there was a significant increase in ^13^C-labeled TCA cycle metabolites and 2HG in high glutamine conditions, indicating that the enhanced astrocytic proinflammatory response in high glutamine conditions is accompanied by increased glutaminolysis for the TCA cycle utilization.

**Figure 6.**
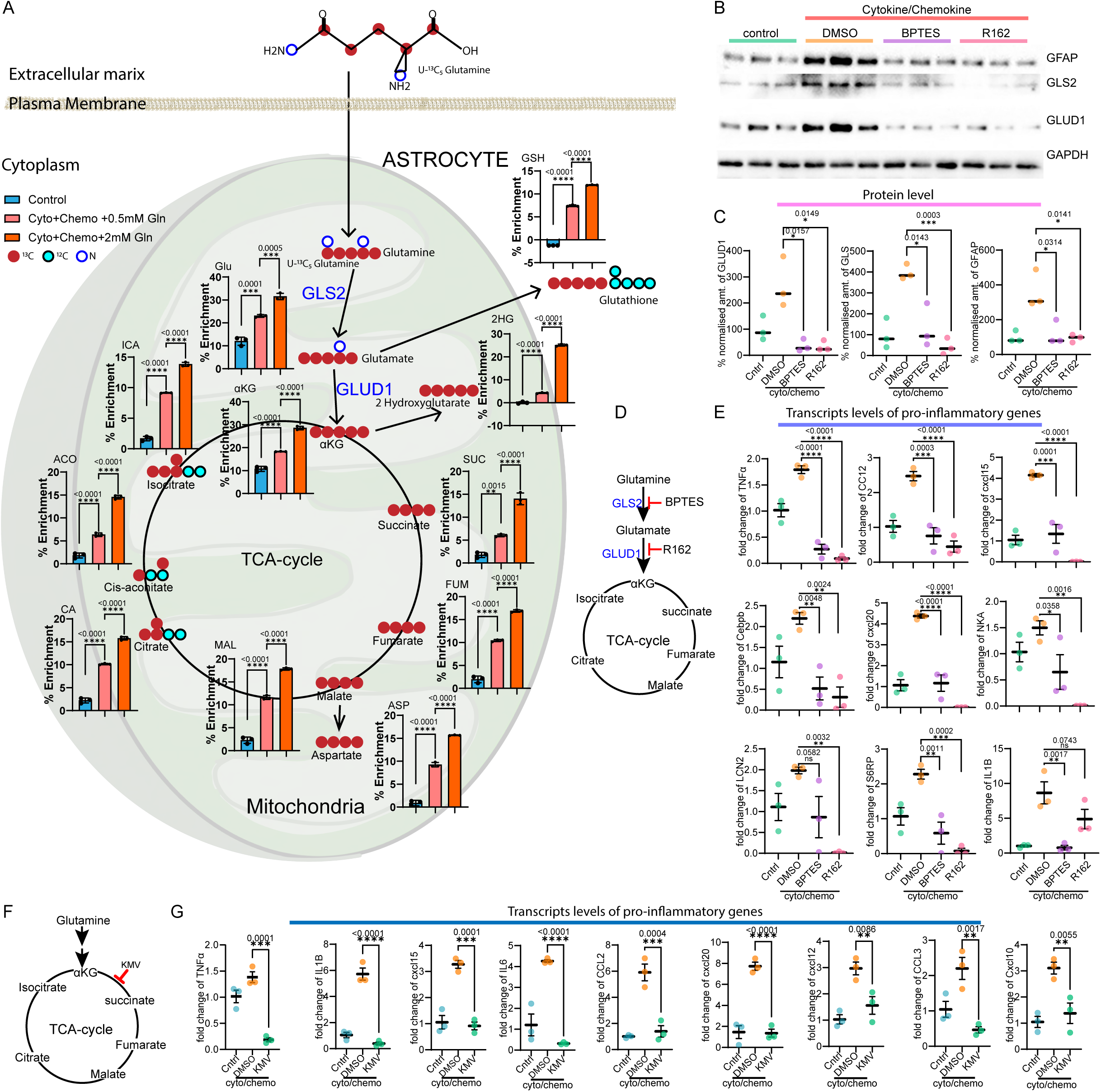
Glutaminolysis fuels reactive astrocytes for a proinflammatory response. (A) U-^13^C_5_-glutamine tracing for glutaminolysis and TCA-cycle metabolites in reactive astrocytes and controls. (B) Immunoblotting for GLS2, GLUD1, and GFAP protein levels in primary reactive astrocytes and following GLS2 or GLUD1 inhibition compared to control. n=3 (C) Densitometric quantification of GLS2, GLUD1, and GFAP protein levels in reactive primary and following GLS2 or GLUD1 inhibition compared to control normalized to GAPDH. n=3 (D) qPCR analysis of proinflammatory gene expression in reactive astrocytes and inhibition of GLS2 or GLUD1 compared to control normalized to GAPDH. n=6. Data significance was calculated using a two-tailed unpaired Student t-test. Error bars represent ± sem. *p<0.5, **<0.005, ***p<0.0005, ****p<0.0001 (E) schematic for TCA cycle block by KMV. (F) qPCR analysis of proinflammatory gene expression in primary astrocytes in physiological glutamine 0.5mM following cytokines and chemokines activation and KMV treatment, normalized to GAPDH. n=3

Previous studies suggested that astrocytes are a key lactate producer, potentially shuttled to neurons^30,31^. To evaluate whether astrocytes utilize glutamine to synthesize lactate, we analyze reactive Ast^C+C^ ^13^C-lactate, finding no significant enrichment under physiological glutamine concentrations compared to control (Figure S6D). However,^13^C-lactate significantly increased following high-glutamine conditions, which needs further investigation. Additional analysis of glycolysis metabolites in reactive Ast^C+C^ revealed no significant changes in fructose-6-phosphate, phosphoenolpyruvate, or pyruvate levels (Figure S6D, S6E). We also did not observe altered pentose-phosphate pathway metabolites D-erythrose 4-phosphate, ribose 5-phosphate, and D-sedoheptulose 7-phosphate (Figure S6F, S6H). However, reactive Ast^C+C^ did increase ^13^C-aspartate levels, an amino acid associated with proinflammatory cytokine IL-1β secretion (Figure 6A)^32^. We further found increased pyrimidine levels in Ast^C+C^ and 2mM Ast^C+C^ (Figure S6G). However, we found no significant changes in purine levels in reactive Ast^C+C^ (Figure S6G).

Next, we determined whether glutaminolysis is necessary for a proinflammatory response and treated reactive Ast^C+C^ with GLS inhibitor (BPTES) or GLUD1 inhibitor (R162)^33,34^. Immunoblotting analysis of reactive Ast^C+C^ revealed a significant increase in glutaminolysis enzymes GLS2/GLUD1 and increased GFAP protein, characteristic of reactive astrocytes in AD (Figure 6B, 6C). Inhibition of GLS2 or GLUD1 significantly decreased glutaminolysis enzymes and GFAP protein levels in reactive Ast^C+C^ (Figure 6B, 6C). We further validated that inhibiting glutaminolysis enzymes GLS2/GLUD1 suppresses proinflammatory gene expression in reactive Ast^C+C^ by qPCR analysis (Figure 6D). This analysis demonstrates that blocking either the conversion of glutamine to glutamate by GLS2 inhibition or the metabolism of glutamate to α-KG by inhibiting GLUD1 decreased proinflammatory gene expression in reactive Ast^C+C^ (Figure 6D, 6E). To determine if TCA cycle utilization of glutamine is necessary for a proinflammatory response, we next treated reactive Ast^C+C^ with alpha-keto-beta-methyl-n-valeric acid (KMV), a structural analog of α-KG that inhibits α-ketoglutarate dehydrogenase complex (KGDHC) activity. KGDHC is an enzyme that converts α-KG to succinyl-CoA in the TCA cycle, an irreversible step. Blocking α-KG conversion to succinyl-CoA substantially reduced several proinflammatory gene expressions of reactive Ast^C+C^ (Figure 6F, 6G). These data suggest that reactive astrocytes increase glutamine uptake for glutaminolysis and TCA-cycle utilization, and blocking glutaminolysis suppresses astrocytic activity.

### Blocking astrocytic glutaminolysis decreases reactive astrocyte and Aβ pathology

To determine whether reactive astrocytes, which increase SNAT2 and glutaminolysis enzymes GLS2/GLUD1, are reliant on glutamine metabolism for a pathological response, we aimed to block glutaminolysis in astrocytes of aged APP/PS1 mice. To target glutaminolysis in astrocytes, we constructed an AAV vector harboring a short CRISPR Cas9-HA tag from *Campylobacter jejuni*^35^ under a truncated GFAP promoter Gfabc1d for selective astrocytic expression. The same construct has single guide RNAs (sgRNA) driven by a U6 promoter targeting the GLUD1 gene (AAV^Glud1/Ko^) or control scramble sequence (AAV^scrmbl^) (Figure 7A). We validated the AAV^Glud1/Ko^ plasmid cleaves the GLUD1 gene in transfected N2A cells by GLUD1-PCR and guide-it-mutation assay, which showed a cleaved DNA with a lower band (Figure 7B). Immunoblotting analysis further demonstrated our AAV^Glud1/Ko^ significantly decreased GLUD1 protein levels compared to AAV^scrmbl^, validating our CRISPR Cas9 GUD1 knockout approach (Figure 7C).

**Figure 7.**
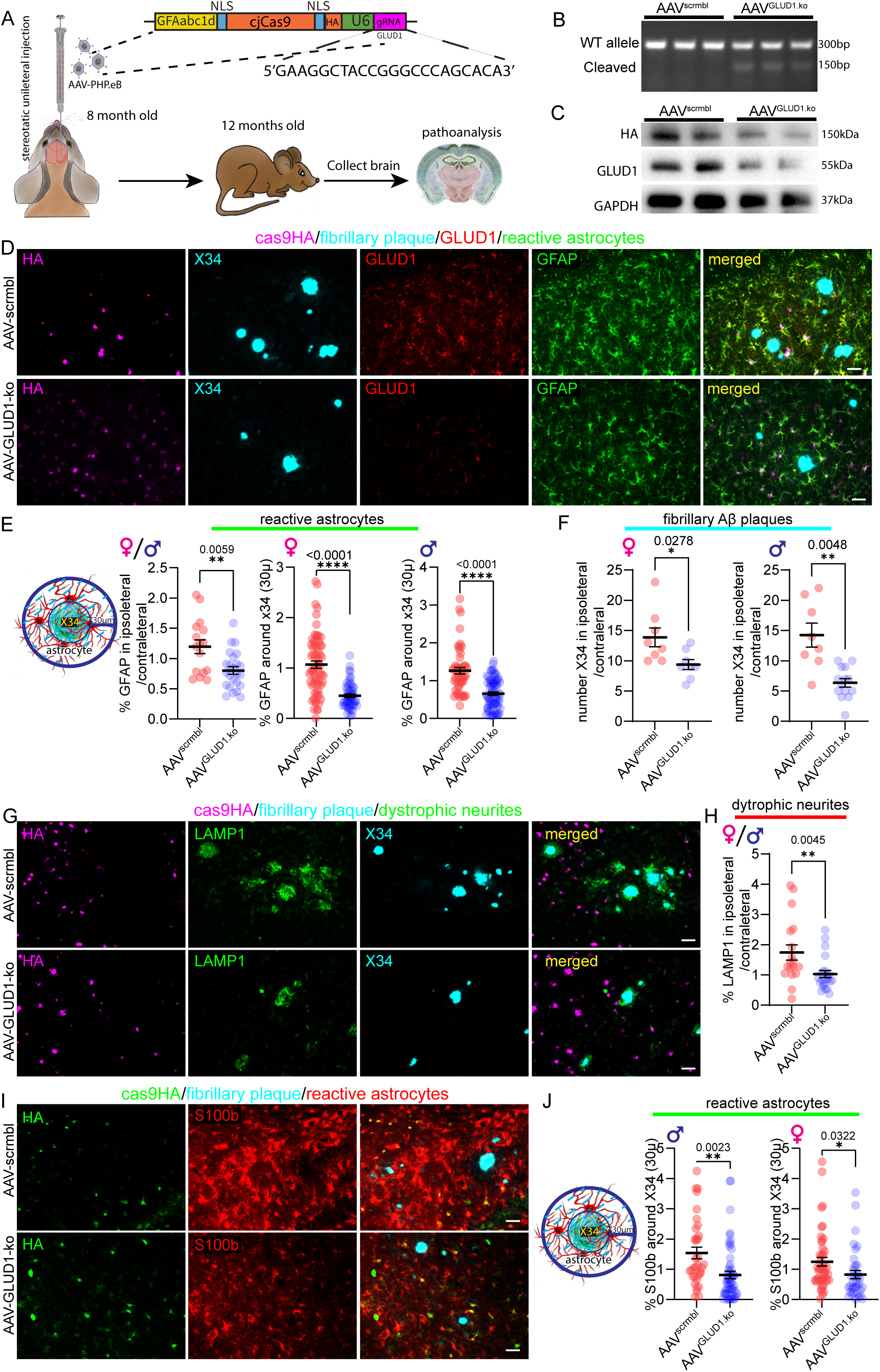
Blocking astrocytic glutaminolysis decreases reactive astrocytes and Aβ pathology. (A) Schematic for timing of AAV-GFAP-Cas9 injections into female/male APP/PS1 mice. (B) Guide-it-Mutation assay demonstrating cleavage of GLUD1 gene with CRISPR-Cas9. (C) Immunoblotting for cjCas9, GLUD1, and gapdh in N2A cells transfected with scramble and GLUD-Ko construct. (D) IF for Cas9-HA, fibrillary plaque, GLUD1, and astrocytes in the hippocampus of APP/PS1-GLUD1 knockout and control. Scale bar, 30μm. (E) Quantification of the percent area of astrocytes around a 30-μm radius of fibrillary plaques in APP/PS1-GLUD1 knockout and control. n= 4-8 (F) Quantification of fibrillar plaque numbers in APP/PS1-GLUD1 knockout and control. n= 4-8 (G) IF for Cas9-HA, dystrophic neurites, and fibrillary plaque in the hippocampus of APP/PS1-GLUD1 knockout and control. Scale bar, 30μm. Scale bar, 30μm. (H) Quantification of the percent area of dystrophic neurites in APP/PS1-NKA knockdown and control. n= 4-8 (I) IF for Cas9-HA and astrocytes in the hippocampus of APP/PS1-GLUD1 knockout and control. Scale bar, 30μm. (J) Quantification of the percent area of astrocytes around a 30-μm radius of fibrillary plaques in APP/PS1-GLUD1 knockout and control. n= 4-8 Data significance was calculated using a two-tailed unpaired Student t-test. Error bars represent ± sem. *p<0.5, **<0.005, ****p<0.0001

We next aimed at blocking astrocytic glutaminolysis, beginning at 8-month-old APP/PS1 when both genders have measurable Aβ pathology and reactive astrocytes. To selectively knock out astrocytic GLUD1, we injected the AAV^scrmbl^ or AAV^Glud1/Ko^ unilaterally into the hippocampus. Following four months post-injection, we analyzed amyloid-related pathologies (Figure 7A). The analysis of astrogliosis revealed a significantly decreased GFAP around the 30μm radius of fibrillary plaques (X34) in both female and male APP/PS1 mice following astrocytic GLUD1 conditional knockout compared to control APP/PS1 mice (Figure 7D, 7E).

We next evaluated the effects of blocking astrocytic glutaminolysis on Aβ pathologies. Initial IF analysis for total Aβ plaques (6E10) revealed no significant change following the conditional astrocytic GLUD1 knockout compared to control APP/PS1 (Figure S7A, S7B). However, IF analysis of fibrillary plaques (X34) associated with neuronal dysfunction displayed a significant decrease in female and male APP/PS1 mice following astrocytic GLUD1 knockout in APP/PS1 mice compared to control APP/PS1 mice (Figure 7D, 7F). To further evaluate a potential neuroprotective effect of blocking astrocytic glutaminolysis, we analyzed dystrophic neurites (LAMP1). This analysis revealed that astrocytic GLUD1 knockout significantly decreased dystrophic neurite accumulation compared to control APP/PS1 mice in both males and females (Figure 7G, 7H).

We further analyzed astrocytic S100β, finding a significant decrease surrounding fibrillary plaques following astrocytic GLUD1 knockout in both male and female APP/PS1 mice compared to control APP/PS1 mice (Figure 7I, 7J). The significant reduction in S100β further suggests astrocytic glutaminolysis is not limited to certain populations of astrocytes, as GFAP is expressed in only a subset of astrocytes. Furthermore, our analyses demonstrate that glutaminolysis regulates reactive astrocytes, and astrocytic GLUD1 knockout suppresses astrocytes’ response to pathology.

## Discussion

Our studies found that reactive astrocytes increase glutamine uptake and glutaminolysis for anaplerosis, a metabolic adaptation that replenishes the TCA cycle metabolites. This novel glutamine metabolic pathway is mediated by increasing the expression of glutamine SNAT2 transporter and glutaminolysis enzymes GLS/GLUD2, which are evident in astrocytes from human AD brain samples, APP/PS1 mouse model of amyloidosis, and primary reactive astrocytes. We further highlight that increasing CNS-glutamine levels with a high dietary glutamine exacerbated reactive astrogliosis and Aβ pathologies in both early and late amyloidosis. Furthermore, we found glutaminolysis sustains astrocytic reactivity through anaplerotic TCA cycle, and blocking glutaminolysis through GLUD1 conditional knockout reduces reactive astrocytes, thereby decreasing Aβ plaques in an *in vivo* amyloid mouse model. In addition, we previously reported that astrocytes increased NKA expression in ALS and AD, which contributed to astrocytic reactivity, but the mechanism was entirely unclear. Our current study further reveals that the NKA is mechanistically interdependent on increased SNAT2-glutamine uptake for astrocyte reactivity. These studies identified astrocytic glutamine metabolism as a key response in pathological conditions, identifying several targetable metabolites and enzymes that could be leveraged to suppress their proinflammatory response.

In the CNS, the flow of glutamine is presumed unidirectional, where astrocytes take up glutamate by excitatory amino acid transporters (EAATs) from synapses for the conversion to glutamine that is resupplied to neurons to generate glutamate for neurotransmission. However, we did not observe an increase in c-fos, a marker for neuronal activity^36^, following the increasing CSF glutamine level through high dietary glutamine as compared to the control diet. This finding suggests that glutamine is not exclusively absorbed by neurons for glutamate-dependent synaptic activity. Rather, glutamine import is higher in reactive astrocytes in pathological conditions, as evident by exacerbated reactivity and pathological responses. Moreover, our study provides a profound understanding, highlighting an increase in SNAT2 expression for glutamine import and glutaminolysis enzymes for glutamine metabolism in reactive astrocytes, thereby defining a new role of glutamine in sustaining astrogliosis and affecting AD pathology independent of neuronal activity, which was previously unknown.

We further show that reactive astrocytes increase their proinflammatory response following increased glutamine concentrations, a characteristic of highly metabolic cells in the periphery^37,38^. Corroborating these results, we demonstrated that increasing CNS-glutamine levels through high dietary glutamine in early disease exacerbates reactive astrocytes, which adversely impact Aβ pathologies, primarily in female mice. As amyloidosis develops more slowly in male APP/PS1 mice, the trending changes in total Aβ plaques potentially indicate that increasing glutamine levels do not affect the disease onset but rather regulate the astrogliosis progression after the disease onset. Consistent with this notion, increasing CNS-glutamine levels after disease onset amplified astrocyte reactivity and Aβ pathology, including dystrophic neurites in both female and male APP/PS1 mice, underscoring the glutamine and astrocytic effects on neuronal health in amyloidosis.

These findings further suggest that dietary glutamine levels, which potentially increase CNS-glutamine, may have a key role in influencing astrocytic responses under pathological conditions. Glutamine is one of the most abundant amino acids in the human diet, and emerging evidence suggests high-protein diets are detrimental, driving chronic diseases, including hypertension and diabetes, known risk factors for neurodegenerative diseases^39–42^. Intriguingly, a protein-restriction diet was reportedly found to slow AD pathology in a mouse model of amyloidosis^43^. Furthermore, a fast-mimicking diet that reduced all macromolecules, including proteins, was also reported to have neuroprotective effects, reducing neuroinflammation in AD mouse models^44^. These findings could be an implication of the lower glutamine diet from a protein-restriction and fast-mimicking diet, which remains to be addressed.

Once inside the cells, glutamine through glutaminolysis replenishes the TCA cycle metabolites. In peripheral inflammatory diseases, glutamine has a key role in inflammation, and blocking the glutamine pathway ameliorates the progression of inflammation ^45,46^. However, the prevailing neuro-centric concept for glutamine utilization has largely hindered further investigation of glutamine’s role in the CNS and neurodegenerative diseases. Yet, our finding of increased glutaminolysis enzymes GLS2/GLUD1 expression in reactive astrocytes of AD human brains and APP/PS1 mice underpins glutaminolysis metabolic adaptation of astrocytes in response to AD pathologies. Although the enzymatic conversion of glutamine to glutamate by GLS2 is a reversible reaction by glutamine synthetase that synthesizes glutamine from glutamate, the increase in GLUD1 enzyme together with^13^C-labeled-α-KG, and TCA-cycle metabolites enrichment indicates a predominant unidirectional glutamine catabolism in reactive astrocytes. Moreover, decreasing astrogliosis with GFAP and S100β and plaque burden after astrocytic GLUD1 conditional knockout supports the unidirectional glutaminolysis sustaining reactivity is not limited to a certain population of astrocytes. These findings indicate that astrocytes indeed utilize glutamine during reactivity, apart from the prevailing neuronal utilization.

The TCA cycle is a fundamental metabolic pathway for energy generation. In addition to energy generation, the TCA cycle functions as a hub for producing several critical cellular metabolites essential for various divergent metabolic pathways, including lipid, amino acid, and nucleotide metabolism. The evidence of increased ^13^C-labeled TCA metabolites suggests that reactive astrocytes employ glutaminolysis as an anaplerotic pathway to replenish TCA metabolites. Our findings of an increased ^13^C-citrate, a crucial substrate for lipid biosynthesis, indicate that glutaminolysis may support the lipid biosynthesis required for hypertrophic morphological changes in reactive astrocytes. It is well known that glutaminolysis provides the amine group required for synthesizing nucleotides. Here, we also observed increased ^13^C-labeled nucleotides—cytosines and uridines—in reactive astrocytes, suggesting that not only does glutamine supply the amine group, but it also provides the carbon backbone for nucleotide biosynthesis, which is necessary for transcriptional changes in reactive astrocytes. Our data further supports that glutaminolysis through the TCA cycle sustains a pro-inflammatory role by contributing to the generation of pro-inflammatory metabolites-2HG, aspartate, and succinate, which are reported to be involved in IL1β production^32,47^. Several reports highlight a positive correlation with pathological astrocytes and increased dystrophic neurites^48,49^. Consistent with the adverse effects of astrogliosis, we found fewer dystrophic neurites in astrocytic GLUD1 knockout APP/PS1, indicating that astrocytic glutaminolysis regulates an inflammatory response detrimental to neuronal health, which may provide strategies for therapeutic intervention.

The limitations of the present study include the exclusive use of the APP/PS1 amyloidosis model, in which reactive astrocytes are also evident in several proteinopathies, including tauopathies. Evaluating other mouse models of proteinopathies could further verify the effect of a high glutamine diet in AD progression. Apart from glutamine as the primary amino acid, SNATs also transport other neutral amino acids with a lesser affinity, which may play a role in astrocyte pathological response. In addition, as a total protein restriction diet appears beneficial for AD, several other amino acids may potentially play an additive role. Our study found that high CNS glutamine levels increased total Aβ production and further increased the dystrophic neurites in a reactive astrocyte-dependent manner.

## Materials and Methods

### Mouse model

The mouse model used in the study was the APP/PSI mouse purchased from the Jackson Laboratory on a C57BL/6J background (Stock 034832-Jax). These mice contain the Swedish mutation in the APP transgene and an L166P mutation in PSEN1, leading to a 3-fold higher APP expression than an endogenous APP mouse. In these mice, the Amyloid beta plaques first appear in the hippocampus in three to four months, followed by the striatum, thalamus, and brainstem in roughly four to five months. We used age-matched APP/PS1-tg and non-tg male and female mice for the experimental research. The IACUC at Washington University School of Medicine approved the study, and it was carried out under the institutional guidelines.

### Primary astrocyte isolation

Primary astrocytes were isolated from P2-P4 CD1 pups (Charles River, 022). After the pups were decapitated, and the skull was opened, and the meninges were stripped, followed by collecting in Ca/Mg-free Hank’s Balanced Salt Solution (HBSS). The cortices collected from 4 pups were washed thrice with HBSS and digested with 2.5% trypsin (GIBCO) in HBSS and 0.2mg/ml DNase (Sigma) at 37 °C for 15 mins. The pellet was washed with HBSS three times, and the tissue was dissociated in HBSS supplemented with 0.4% mg/ml DNase and spun at 1500 rpm for 5 mins. The supernatant containing cell debris was discarded, and the pellet was resuspended in DMEM media supplemented with 10% FBS and 1x PenStrep (DMEM-FP) and seeded in a 10cm petri dish. After 24 hrs of incubation at 37 °C and 5% humidity, the media from the culture plate is discarded and replaced with fresh DMEM-FP and cultured for 7 days. After another 7 days, astrocytes were passaged into two 10cm petri dishes coated with 1: 400 dilution of Geltrex (GIBCO) for overnight in DMEM-FP media.

### Immunoblots

To calculate the amount of total protein needed from APP/PS1 mice and humans, we homogenized cortical brain samples with a pellet pestle in RIPA Buffer at a concentration of 10mg/400.0ul (50Mm Tris base, 150mM NaCl, Na-deoxycholate, 1% Triton, 0.1 SDS, 5mM EDTA, 1mM Na3UO4, cocktail of protease and phosphatase inhibitors). Next, utilizing a probe sonicator, the samples were sonicated on ice for one minute at 60% amplitude (1 sec on, 1 sec off intervals). Following sonication, protein concentration was determined using a BCA kit (Thermo Fisher), and equal quantities from each sample were solubilized in an SDS-loading buffer. Samples were then loaded on a 10% gel and transferred onto a nitrocellulose membrane, which was then blocked with 3% BSA in TBS-T buffer for an hour and incubated in primary antibodies GLUD1 (1:4000), GLS2 (1:4000), GFAP (1:15000), SNAT2 (1:4000), NKA (1:16000), LCN2 (1:500), GAPDH (1:5000). Following three TBS-T washes, the secondary antibody was applied (1:10000) for 1 hour at room temperature (RT) and further washed with TBST (10 min. each). The immunoblots were then developed and analyzed using Sigma Aldrich ECL and the Bio-Rad ChemiDoc gel imaging system. Densitometric analysis was conducted using image J software.

### Intracranial hippocampus injection of injection of AAV

AAV-GFAP-cre, AAV-GFAP-GFP were purchased from (UNC). AAV-GFAP-cjCas9-GLUD1Ko was first generated in pX601-CMV-SaCas9(reopt)-3xHA-U6-BsaI(sgRNA).gbk^50^ backbone replacing CMV with Gfabc1d promoter (Age1/Spe1) and SaCas9 with cjCas9 from pAAV-EFS-CjCas9-eGFP-HIF1a^35^ (Age1/EcoR1). The Glud1 guide RNA and scramble RNA oligos were ligated into the Bsa1 site. AAV viruses were generated following previously described^51^. We injected 8 month old APP/PS1 mice with 2μl of 1×10^12^ AAV in the dentate gyrus (bregma; −2.5mm, lateral; 2.0mm, depth; −2.0mm) at 0.2μl/min. The injected brains were collected at 12 months.

### High Glutamine supplementation in APP/PS1 mice

APP/PS1 mice were aged to 6 months for the young cohort and 12 months for the old cohort. They were gavaged daily with either water as control or Glutamine solution in water (1 gm/kg of body weight) daily for 60 days with 22-gauge and 1.5-inch gavage needle.

### Brain collection and preparation

At the designated time of the endpoint, mice were anaesthetized with isoflurane to collect their brains through perfusion with chilled 1xPBS, followed by brain dissection and immersion in 4% PFA. Twenty-four hours later, perfused mice brains were changed from PFA to 30% Sucrose in 1x PBS. Brains were subsequently cut into 30-micron sections using a microtome (Leica) and then preserved in a 24-well plate with cryoprotectant (components of cryoprotectant) and stored at −20°C.

### Immunofluorescence

Immunofluorescence staining was performed using GFAP(1:8000), GLS2 (1:2000), GLUD1 (1:2000), S100B (1:4000), IBA1(1:2000), 6E10(1:2000), cFos (1:10000), Vimentin (1:5000), SNAT1 (1:2000), SNAT2 (1:2000), MAP2 (1:5000), HA (1:4000), NKA (1:10000), LAMP1 (1:4000), and X34 (1:5000). Free-floating sections were incubated in Sudan Black (Sigma)(100mg/100ml in 70% EtOH) for 15 minutes to quench lipofuscin and then washed with 0.02% Tween-X-100 in 1xPBS for 5 minutes three times. Next, sections were incubated in 40% ethanol and 1xPBS with X34 (cat number, 1:5000), NaOH (12N, 1:500) for 20 minutes and washed with 40% ethanol + 1XPBS for 3 times 2 minutes each and further washed with 1xPBS for 10 minutes. Sections were then permeabilized in 0.3% Triton in 1xPBS for 10 minutes, transferred to 1x PBS for 5 minutes, and blocked in 3% bovine serum albumin, 0.1% Triton X-100 in 1xPBS for 1 hour and then incubated with primary antibody diluted in blocking buffer overnight at 4°C. The next day, sections were incubated with secondary antibodies diluted in blocking buffer for 2 hours at RT and washed in PBS for 20 minutes three times. Once washed, the sections were mounted and sealed with Fluoromount-G (Thermo Fischer, 00-4959-52) and stored in the dark at 4°C.

### Immunofluorescent staining of Human tissue

Human AD patients’ and control clinical diagnostic criteria and validation were performed as previously described^52,53^. Parafilm-embedded tissues were dewaxed in 100% xylene for 15 mins each for 3 times, followed by 5 mins in xylene/ethanol (1:1 ratio). The tissues were washed in ethanol 100%, 95%,75% and water sequentially two each time for 2 mins. Antigen retrieval of the dewaxed tissues was done in pre-heated (95°C) sodium citrate (10 mM) + 0.05% Tween-20 for 30 mins, followed by cooling down for 20 mins at RT and rinsing with 0.02% Tween-20 prepared in 1XPBS twice for 2 mins each. The dewaxed-antigen retrieved human tissues were immunostained following the protocol mentioned in the Immunofluorescent section.

### Image acquisition and analysis

Images were acquired using a 20X magnification Keyence BZ8100 scanner for % area coverage for pathology analysis and stitched using a BZ-X800 analyzer. Images for analysis of patho-astrocytes around the fibrillary plaques were acquired at 40X magnification using a Keyence BZ8100 scanner at different regions (CA1, CA3, dentate gyrus, and piriform cortex). Representative images for SNAT2, GLUD1, and GLS2 in APP/PS1 brain and human brain were acquired at 100x magnification with a z-stack of 0.5μm pitch and stitched with the BZ-X800 analyzer. All immunofluorescent analyses were performed using Image J software (NIH). Analysis of X34, 6E10, IBA1, and LAMP1 was done by tracing the different regions of the brain, and similar thresholds were applied to all the images, and percent area coverage was measured using Image J. For analysis of GFAP and S100β around the X34 plaques were done by 30μm around X34 was analyzed, similar thresholds were applied, and the percent area coverage was measured using Image J. measurement of c-Fos-expressing neurons was conducted using image J, measuring the count in the hippocampus and cortex.

### Cerebrospinal fluid extraction

APP/PS1 wildtype mice were gavaged for 7 days with high Glutamine (10g/Kg). On day 8, mice were gavaged for 45 mins and anesthetized with 40mg/kg of Na-pentobarbital (Fetal plus). mice head was secured using ear bars. A vertical incision of a length similar to the ears along the anterior-posterior axis between the ears was made using sharp scissors, followed by cutting the layer of fascia and muscle. The muscles in the posterior neck, right below the skull were pulled apart from each side using retractors to expose the dura. Once the dura is exposed, we located cisterna magna and a sterile pulled-capillary tube with a bevel tip connected to a 3-stopper valve was inserted in the dura membrane right above the cisterna magna and allowed the CSF to flow along the tube by using negative pressure. The CSF was collected in a fresh tube and stored at −80° C.

### Measuring Glutamine from CSF and brain lysate

Glutamine concentration in the human brain lysate and mouse CSF was measured using Glutamine/Glutamate Glo Assay (Promega, J8021). Briefly, 4-5mg of human cortical brain was sliced and lysed using 1.125 ml of homogenization buffer (mixture of 50 mM Tris, pH 7.5, 0.6 M HCl in a 1:8 ratio). After homogenization using tissue tearor for 20-30 sec, 0.125 ml of 600mM Tris (pH 8.5) was followed by the manufacturer’s protocols. Luminescence was recorded using a Varioskan-Lux Multimode Microplate reader (Thermofisher Scientific). Glutamine concentration in the CSF was measured after deprotonation using 0.6N HCl and neutralization using 600mM Tris (pH 8.5) and following the manufacturer’s instructions.

### Glutamine rhodamine assay

Primary astrocytes were seeded in a coverslip coated with geltrex overnight. After culturing for 5 days astrocytes were treated with chemokine and cytokine cocktail as mentioned above, along with 1.2μM of digoxin (Sigma Aldrich D-029) and 3 mM of MeaIB (Sigma Aldrich, M2383) in separate groups for 48 hours. After 48 hours, astrocytes were treated with 0.5 mM of rhodamine-B as negative control and 0.5 mM of rhodamine-labelled glutamine for 10 mins. Astrocytes were fixed with 4% PFA for 10 minutes, followed by permeabilization with 0.3% Triton X-100 in PBS for 10 mins. Astrocytes were immunostained with GFAP and the coverslips with astrocytes were mounted on slides using Fluoromount-G with DAPI (ThermoFisher Scientific, 00-4959-52).

### Generation of NFκB reporter lentivirus primary astrocytes transduction

NFκB-reporter plasmid was generated using NFκB minimal response element sequence obtained from ^54^and cloned in pLL3.7 vector ^55^ by replacing CMV promoter sequence using Not1 and Age1 restriction enzymes. 15-18 million 293T cells were seeded in a 15cm tissue culture plate for 16-20 hrs in DMEM supplemented with 5% fetal bovine serum, 0.1% penicillin-streptomycin, and 0.1% MEM Non-essential Amino acids solution (Gibco, 11140050) and transfected with NFκB reporter plasmid, pMD2.G (Addgene, 12259) and psPAX2 (Addgene, 12260) using polyethylenimine (PEI) prepared as mentioned previously. Fresh media was replaced after 3 hrs of transfection and incubated for 3 days. Media containing lentivirus were collected and centrifuged at 1500 rpm for 5 mins, then filtered with a 0.45μm Millex PVDF syringe filter (Sigma, SLHVR33RS). Lentivirus was collected by spinning at 25000 rpm for 2 hrs in an Ultracentrifuge (ThermoFisher, Sorvall Wx100+ Ultracentrifuge). Media were discarded and viral pellets were dissolved in PBS for 1 hour at RT and stored at −80°C.

### RNA isolation for qPCR

2×10^5^ primary astrocytes were cultured in 6-well plates for 5 days in Neurobasal media, followed by treatment with TNFα+IL1β+IL6 (50ng/ml,1ng/ml,10ng/ml) for 48hrs. Cells were washed with 1XPBS, and 1 mL of TRIzol was added to the cells, followed by RNA isolation, after adding 200μl of chloroform and centrifugation at 12000g for 15 mins. The aqueous layer was collected in fresh tubes, followed by adding 500μl of isopropanol and mixing 10 times. The samples were incubated in −20 for an hr and centrifuged at 12000g for 15 minutes. The supernatant was discarded, and RNA pellets were washed with 80% ethanol and centrifuged for 10 minutes at 12,000g. DNAse treatment and reverse transcription were performed following the manufacturer’s protocol of High-Capacity cDNA Reverse Transcription Kit (Thermofisher,4368841). qPCRs were conducted using the manufacturer’s protocol of Sybr Green Universal Master Mix (ThermoFisher, 4309155) in Quant-Studio by Applied Biosystem.

### Glutamine ^13^C tracing

Primary astrocytes were plated in 35mm plates for 5 days in Neurobasal media. Cyto/chemo astrocytes were treated with a cytokine/chemokine cocktail using ^13^C glutamine for 48 hrs. Cells were collected in methanol and metabolic analysis was conducted using LC/MS-MS following the previously described^56^.

### Statistical analysis

All results were reported as means± SEM, and all statistical analyses were conducted in Prism 10 (GraphPad). Significant differences between experimental groups were performed with unpaired Student’s two-tailed *t* test or One-way ANOVA, and the Mann-Whitney post hoc test was used for assessing significance between more than two groups or more. p < 0.05 was considered a statistically significant difference. \**p* < 0.05, \*\**p* < 0.01, \*\*\**p* < 0.001, and \*\*\*\**p* < 0.0001. Each exact *p-*value is mentioned in each respective graph. The value of *n* per group is indicated within each figure legend.

## Acknowledgement

We thank the staff of the Division of Comparative Medicine (DCM), Washington University, School of Medicine. We thank the patients who participated in this research, their families, and the investigators and staff at Washington University, School of Medicine, Knight Alzheimer’s Disease Research Center (ADRC). This work was supported by the Hope Center Viral Vectors Core at Washington University School of Medicine. We acknowledge Dr. Philip Lorenzi and Lin Tan in the Metabolomics Facility, supported by The University of Texas MD Anderson Cancer Center and funded by P30CA016672, for ^13^C glutamine tracing mass-spec and data analysis.

## Author contributions

S.S.D. and G.G. conceived and designed the study. S.S.D. performed all experimental aspects, including stereotactic injections, immunoblots, histopathological staining and analysis, high-glutamine diet analysis, and qPCR analysis, cloned all constructs, and generated viruses for unilateral brain injections. S.L.H. performed tissue harvesting, brain tissue dissections, histopathological staining, and mouse husbandry. I.L. provided the rhodamine-labeled glutamine. S.S.D. and G.G. wrote the manuscript with input from all authors. All authors approved the manuscript.

## Supplementary Figure legends

**Figure Supplementary 1.**
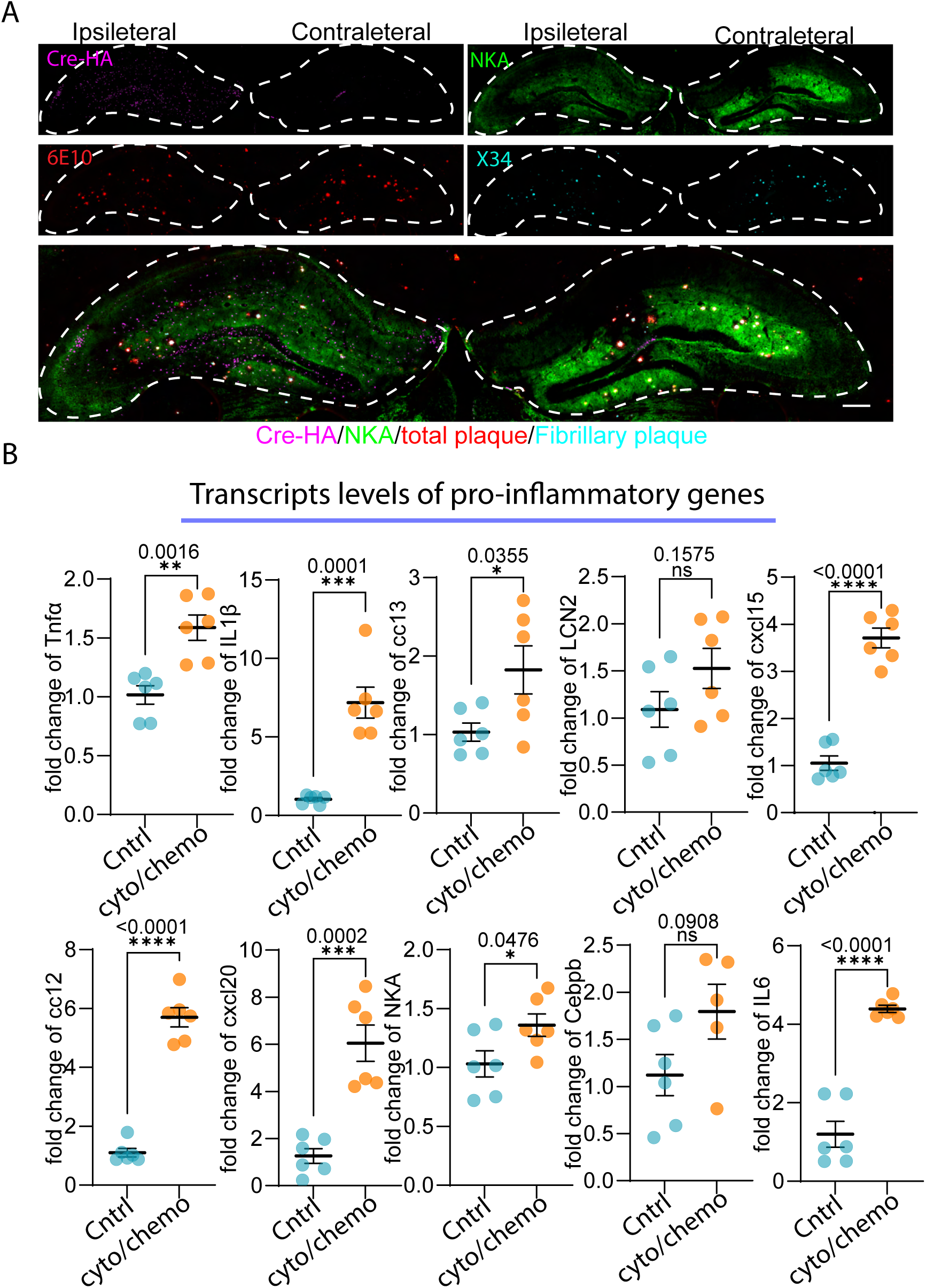
Astrocytic NKA knockdown in aged APP/PS1 mice and proinflammatory gene analysis of reactive astrocytes. (A) IF for Cre-HA, NKA, fibrillary and total plaques in whole hippocampus of APP/PS1-NKA knockdown and control. Scale bar, μm. (B) qPCR analysis of proinflammatory gene expression in primary astrocytes and reactive astrocytes with cytokines and chemokines normalized to GAPDH. n=6

**Figure Supplementary 2.**
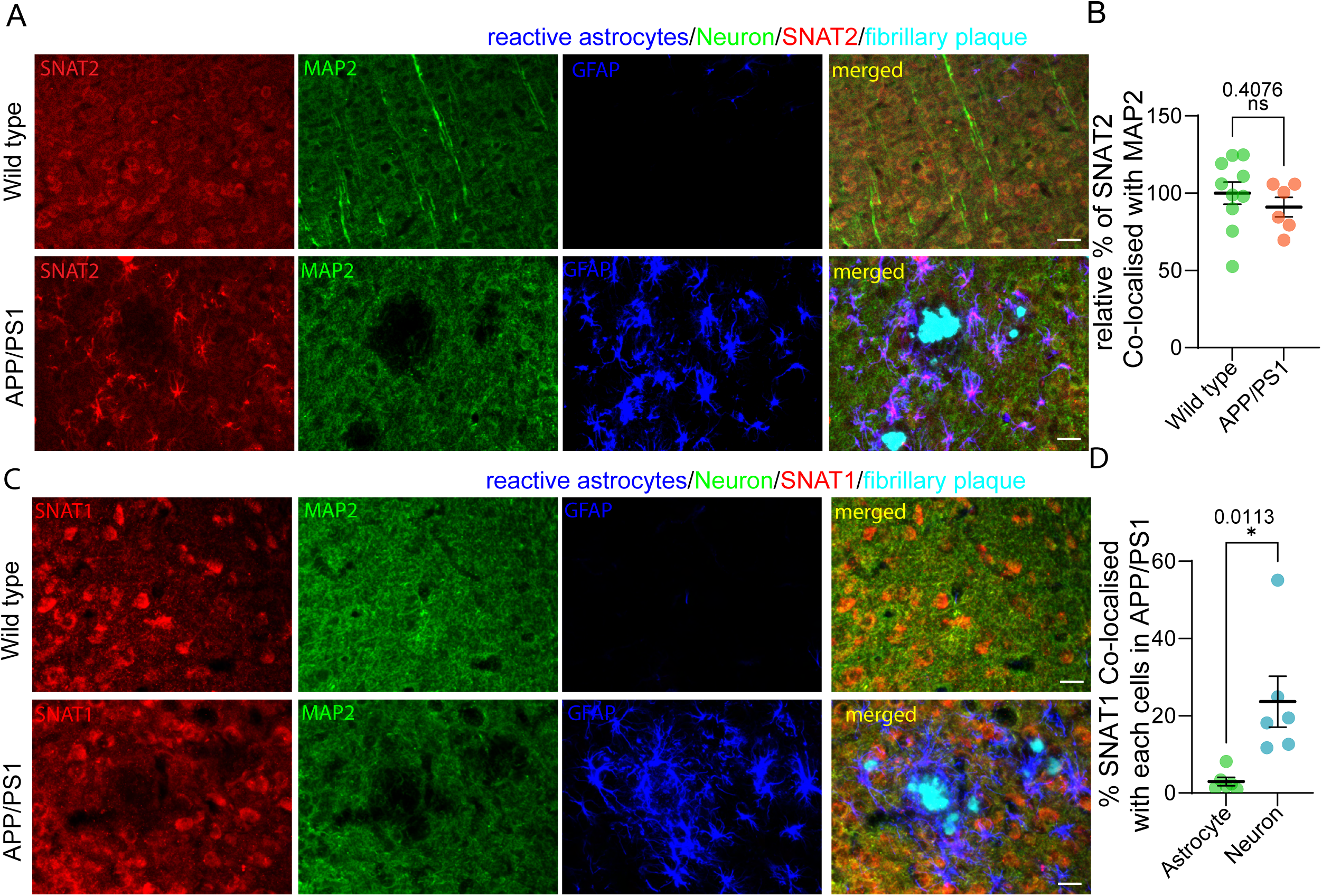
SNATs expression in aged APP/PS1 mice. (A) IF for astrocytes, MAP2, SNAT2, and fibrillary plaques in APP/PS1 and control wild type mice. n=3-5 Scale bars 30μm. (B) Quantification of % area coverage of SNAT2 colocalized with neurons in APP/PS1 mice compared to wild-type mice. n=3.(C) IF for astrocytes, MAP2, SNAT1, and fibrillary plaques in APP/PS1 and control wild type mice. n=3 Scale bars 30μm (D) Quantification of % area coverage of SNAT1 colocalized with astrocytes and neurons in APP/PS1 mice. n=3

**Figure Supplementary 3.**
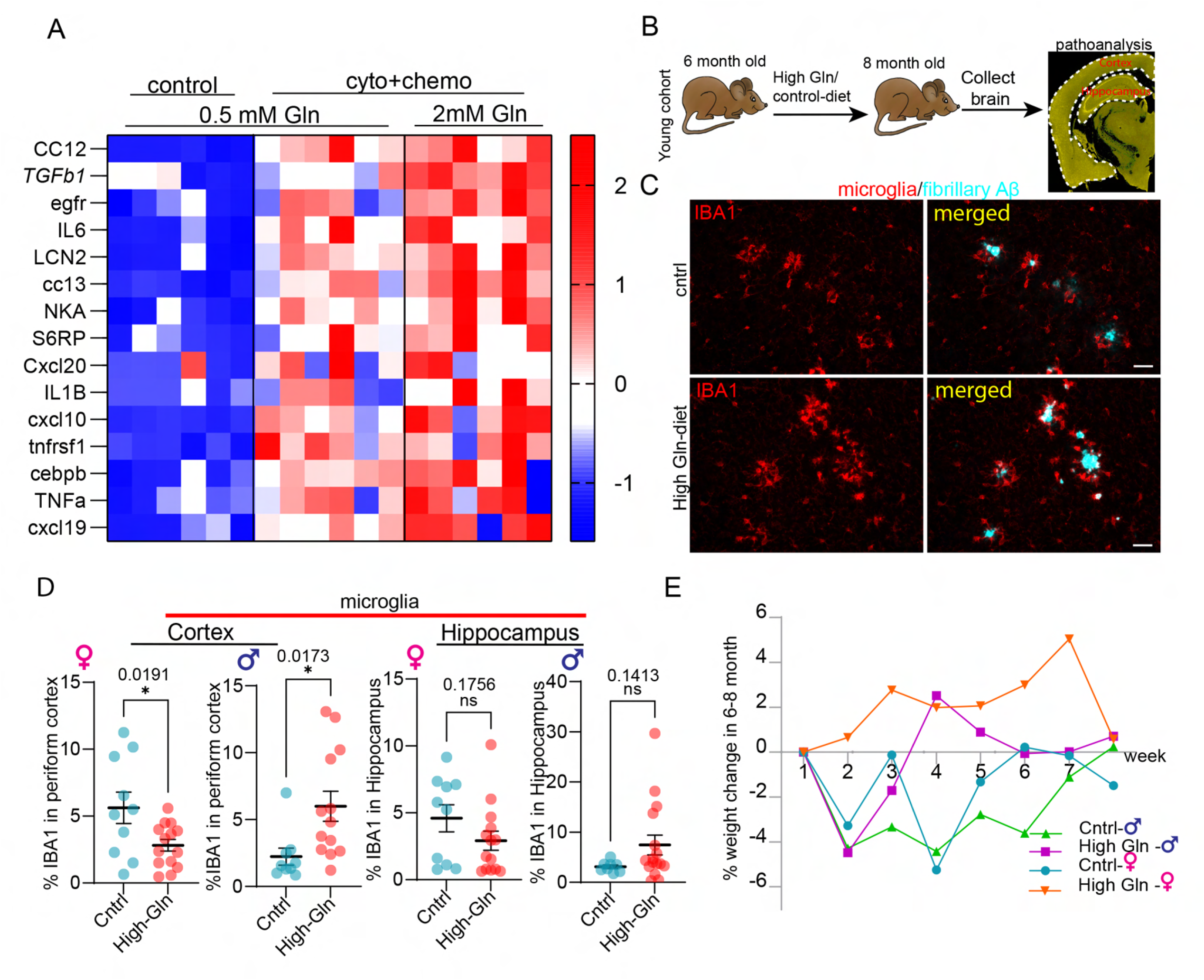
Increasing glutamine levels exacerbate reactive astrocytes *in vitro* and *in vivo* before disease onset in APP/PS1 mice. (A) qPCR analysis of proinflammatory gene expression in primary astrocytes in physiological glutamine 0.5mM and high 2mM conditions following cytokines and chemokines activation, normalized to GAPDH. n=6 (B) Schematic of a high-glutamine diet timeline before overt amyloidosis in APP/PS1 mice. (C) IF for microglia and fibrillary plaques following a high-glutamine diet or a control diet in APP/PS1 mice. Scale bars 30μm. (D) Quantification of the percent area of microglia in the cortex/hippocampus following a high-glutamine diet compared to control diet in APP/PS1 mice. n= 5-7. (E) Weight analysis of APP/PS1 mice following a high-glutamine diet or control diet. n= 5-7

**Figure Supplementary 4.**
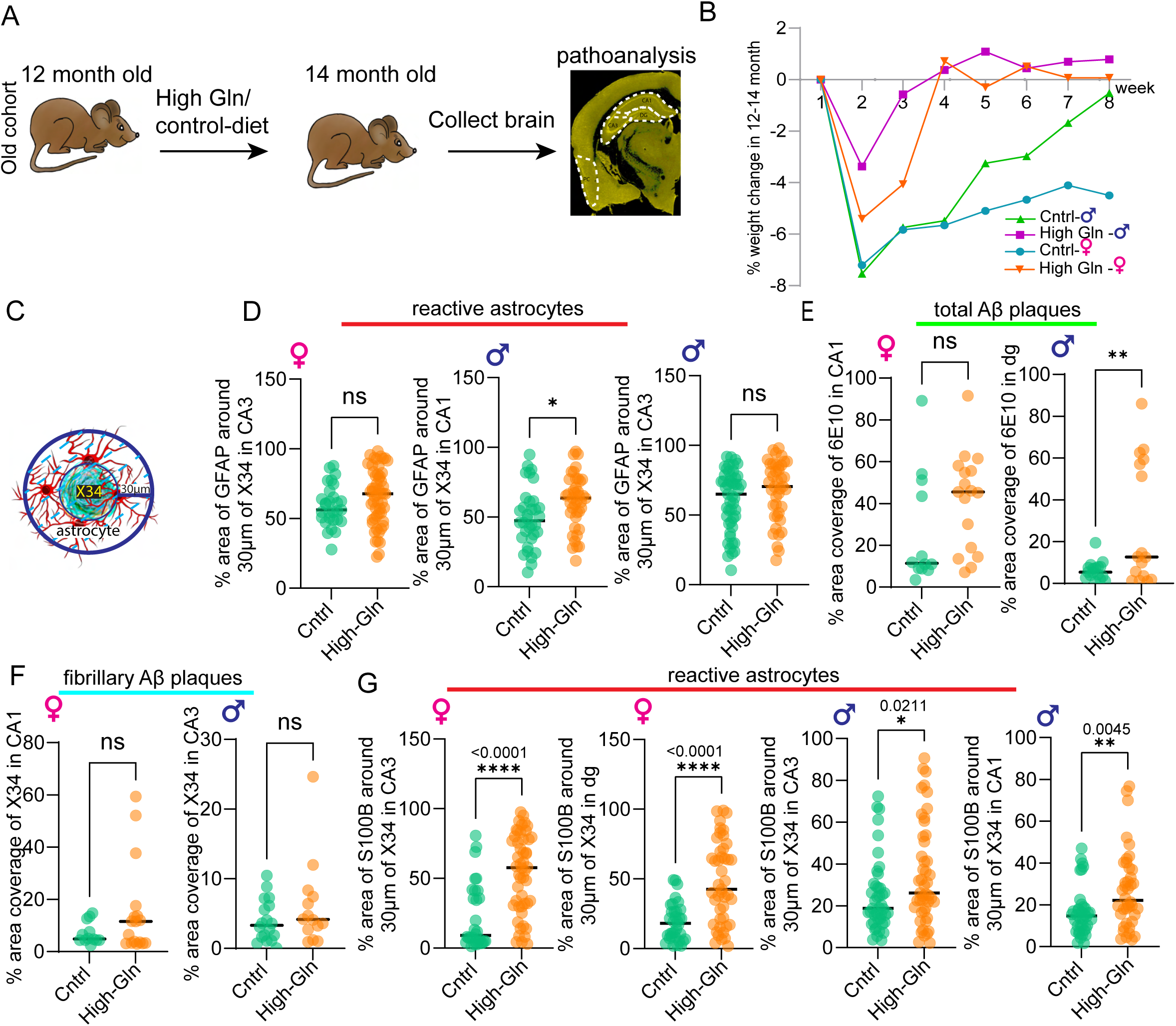
Increasing CNS-glutamine levels after disease onset exacerbates reactive astrocytes and Aβ pathologies. (A) Weight analysis of APP/PS1 mice following a high-glutamine diet or control diet. n=6-9. (B) Quantification of the percent area of astrocytes around a 30-μm radius of fibrillary plaques following a high-glutamine diet compared to control diet. n= 6-9. (D) Quantification of the percent area covered by total plaques in the cortex/hippocampus following a high-glutamine diet compared to control diet. n=6-9. (E) Quantification of the percent area covered by fibrillary plaques in the cortex/hippocampus following a high-glutamine diet compared to control diet. n= 6-9. (F) Quantification of the percent area of astrocytes around a 30-μm radius of fibrillary plaques following a high-glutamine diet compared to control diet. n= 6-9

**Figure Supplementary 5.**
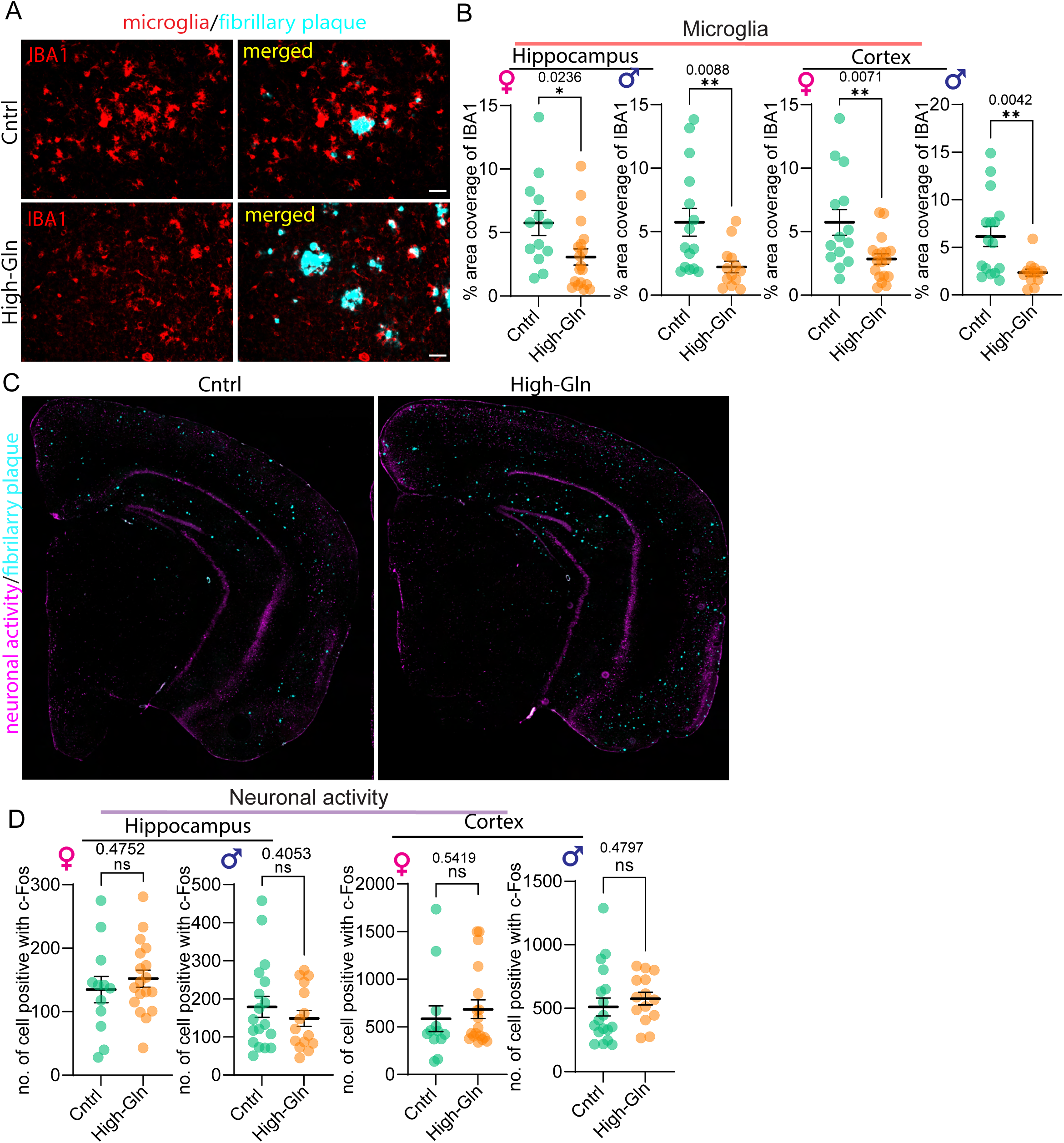
Increasing CNS-glutamine levels alter microgliosis. (A) IF for microglia and fibrillary plaques of APP/PS1 mice following a high-glutamine diet or control diet. n= 6-9. (B) Quantification of the percent area covered by microglial in the cortex/hippocampus following a high-glutamine diet compared to control diet. n= 6-9 (C) IF for neuronal activity and fibrillary plaques of APP/PS1 mice following a high-glutamine diet or control diet. n= 6-9. (D) Quantification for the number of c-FOS positive cells in APP/PS1 mice following a high-glutamine diet or control diet. n= 6-9

**Supplementary Figure 6.**
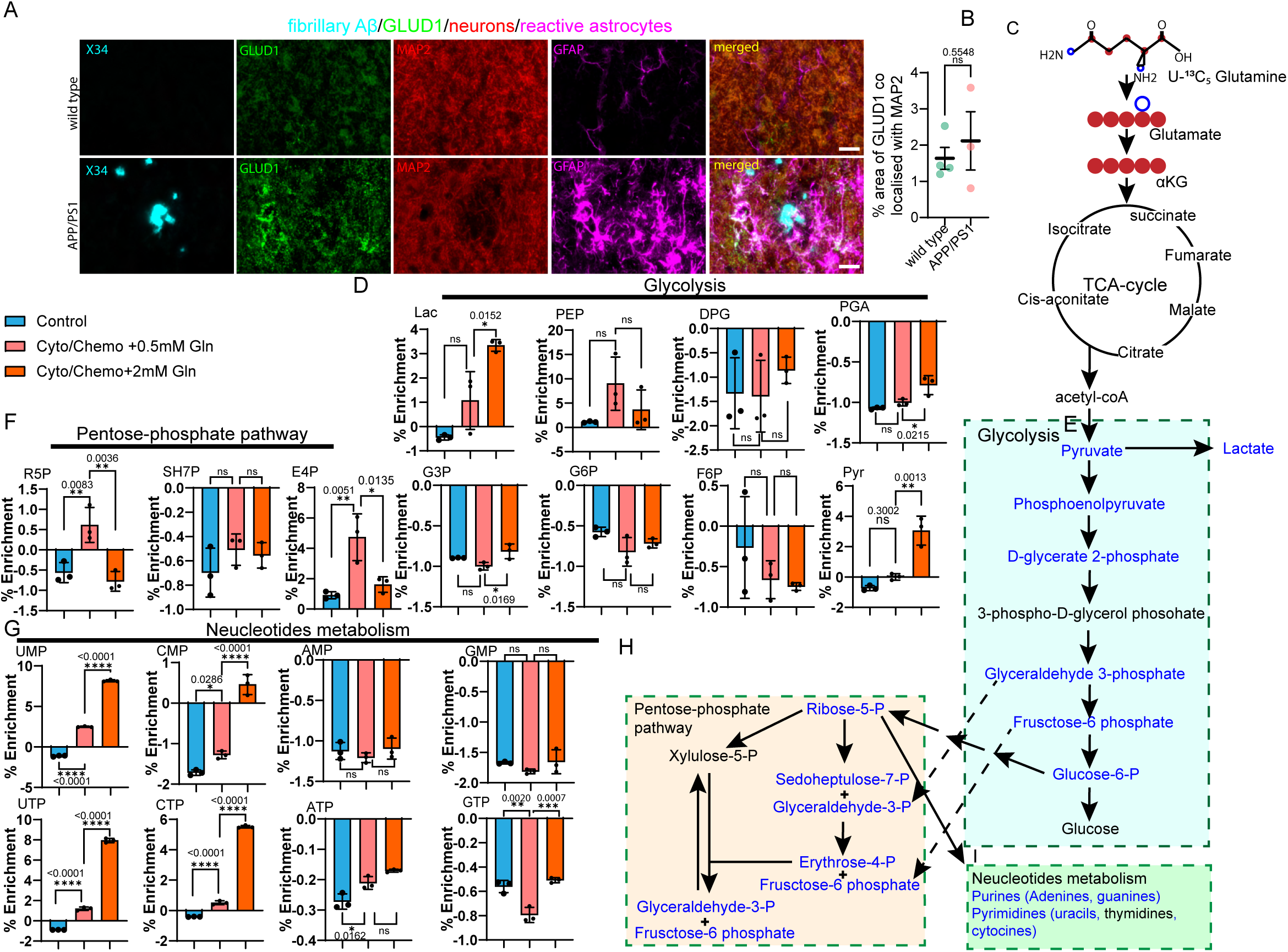
Glutaminolysis GLUD1 in APP/PS1 mice and ^13^C_5_-glutamine tracing of reactive astrocytes. (A) IF for GLUD1, fibrillary plaques, neurons, and astrocytes in APP/PS1 and wild-type mice. Scale bars 30μm (B) Quantification of percent area coverage of GLUD1 colocalized with astrocytes and neurons in APP/PS1 mice compared to wild-type mice. n=3-4. (C) Schematic for TCA cycle metabolites. (D) U-^13^C_5_-glutamine tracing for glycolysis in reactive astrocytes and controls. n=3. (E) Schematic for Glycolysis metabolism (F) U-^13^C_5_-glutamine tracing for the pentose phosphate pathway in reactive astrocytes and controls. n=3. (G) U-^13^C_5_-glutamine tracing for nucleotide metabolism in reactive astrocytes and controls. n=3. (H) Schematic for the Pentose-phosphate pathway. (I) Schematic for nucleotide metabolism.

**Supplementary Figure 7.**
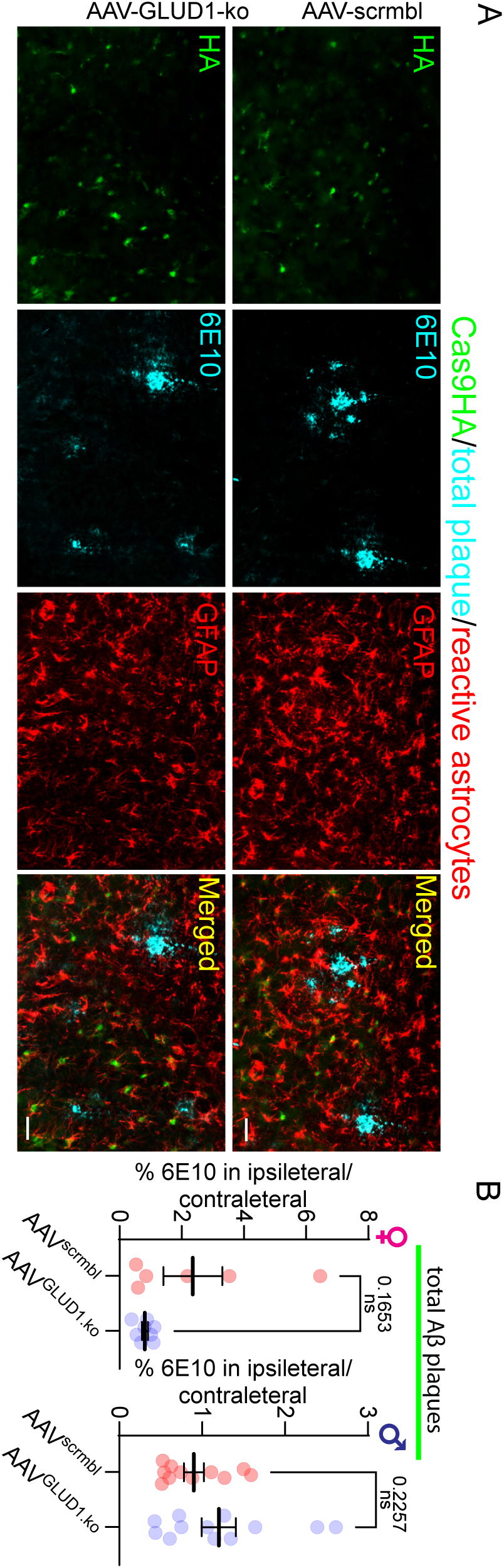
Blocking astrocytic glutaminolysis decreases reactive astrocytes and Aβ pathology. (A) IF for Cas9-HA, total plaques, and astrocytes in the hippocampus of APP/PS1-GLUD1 knockout and control. Scale bar, 30μm. (B) Quantification of the percent area of total plaque in APP/PS1-GLUD1 knockout and control. n= 4-8

